# Reverse gyrase and 3D genome architecture suppress hyperthermophile genome instability arising from horizontal gene transfer

**DOI:** 10.64898/2025.12.08.692931

**Authors:** Kodai Yamaura, Naomichi Takemata, Takeshi Yamagami, Yoshizumi Ishino, Haruyuki Atomi

## Abstract

Reverse gyrase (Rgy), a distinctive topoisomerase conserved in all hyperthermophiles, has the unique ability to introduce positive DNA supercoils. It has long been hypothesized that Rgy overwinds genomic DNA to prevent its detrimental denaturation at high temperature. However, its role *in vivo* has remained unresolved for more than four decades. In the course of investigating how Rgy affects genome organization in the archaeon *Thermococcus kodakarensis*, we find that Rgy suppresses heat-induced clustering of AT-rich genes, most of which bear signatures of horizontal gene transfer. This function depends on the topoisomerase active site of Rgy, underscoring a critical role of its action on DNA topology. The clustering of AT-rich genes is accompanied by their aberrant recruitment of the single-stranded DNA-binding protein RPA, indicative of extensive DNA melting. Genetic analysis further provides evidence that RPA sequesters denatured loci and mitigates genome instability in the absence of Rgy. We propose that DNA topology and higher-order genome organization constitute a multilayered mechanism that stabilizes horizontally acquired genes at high temperature.

## Introduction

The double helical structure of DNA can adopt various topological forms—relaxed, supercoiled, knotted, and unknotted states. Regulated interconversion between these forms is critical to carry out DNA transactions. To modulate DNA topology, cells use a variety of DNA topoisomerases, which catalyze transient breakage of DNA to alter its topological state^1, 2^. Topoisomerases can be classified into two types: type I topoisomerases cleave one strand of double-stranded DNA (dsDNA), while type II topoisomerases cut both strands. Type I and type II enzymes can further be divided into three (IA, IB, and IC) and two (IIA and IIB) subtypes respectively according to structural and mechanistic differences. Due to the essential roles of topoisomerases, as well as the potentially detrimental DNA cleavage reactions that they catalyze, functional defects in topoisomerases can easily lead to cell death and various diseases.

Reverse gyrase (Rgy) is a type IA topoisomerase that relaxes negatively supercoiled DNA and further introduces positive supercoils, whereas the other topoisomerases generate relaxed or negatively supercoiled DNA^3^. This positive supercoiling activity is carried out through functional coupling of Rgy’s ATP-dependent N-terminal superfamily 2 helicase domain and C-terminal topoisomerase domain^4, 5, 6, 7, 8, 9^. An early study observed the coincidence of Rgy and a positively supercoiled plasmid in the thermophilic archaeon *Sulfolobus acidocaldarius*^10^. This finding has led to the hypothesis that Rgy-mediated introduction of positive supercoiling (DNA overwinding) enhances DNA stability at high temperature by preventing thermal denaturation of dsDNA and subsequent breakage of fragile single-stranded DNA (ssDNA). Comparative genomics showed that Rgy is present almost exclusively in hyperthermophiles (optimal growth temperature: ≥ 80°C) and extreme thermophiles (optimal growth temperature: ≥ 65°C)^11, 12^. Genetic studies further revealed that loss of Rgy causes severe growth defects in the hyperthermophilic euryarchaea *Thermococcus kodakarensis* and *Pyrococcus furiosus* and the extremely thermophilic crenarchaeon *Saccharolobus islandicus*^13, 14, 15^. In the former two cases, growth inhibition has been shown to increase with environmental temperature and become lethal at around 95°C. The data from genomics and genetics support the notion that Rgy-mediated control of DNA topology is critical for the genome integrity in hot environments, especially at growth temperatures of hyperthermophiles.

Despite the widely accepted role of Rgy for hyperthermophilic life, there is surprisingly no direct *in vivo* evidence that the positive supercoiling activity of Rgy prevents genomic DNA from melting. Moreover, plasmid DNA in thermophilic cells can be both positively and negatively supercoiled, raising the possibility that DNA positive supercoiling is not crucial for its thermal stability *in vivo*^16, 17, 18^. In line with the dispensability of positive supercoiling, certain hyperthermophiles can endogenously or exogenously express DNA gyrase, a prokaryotic type IIA topoisomerase that removes DNA positive supercoils and instead introduces negative supercoils^19, 20^. *In vitro* studies have proposed alternative mechanisms of how Rgy increases the fitness of thermophiles. For instance, biochemical analyses have suggested that Rgy can act as a heat-protective DNA chaperone independently of the supercoiling activity^21^. Rgy can also destabilize Holliday junctions and inhibit DNA synthesis of translesion DNA polymerase *in vitro*, hinting at its potential roles in DNA repair^22, 23, 24^. Given these findings, the precise role of Rgy *in vivo* remains to be elucidated^25^.

Recent advances in chromosome conformation capture (3C) technologies (Hi-C, 3C-seq, Micro-C, etc.) and microscopy have enriched our understanding of how the DNA double helix is organized into higher-order structure in the three domains of life^26, 27, 28^. A variety of factors, such as Structural Maintenance of Chromosomes (SMC) complexes and DNA-protein condensates, affect the 3D genome organization and thereby control genomic functions^27, 29, 30^.

Topoisomerase-mediated regulation of DNA topology also impacts the higher-order genome folding. For instance, a Hi-C study has revealed that inactivation of *Escherichia coli* Topoisomerase IV (Topo IV), a type IIA topoisomerase/decatenase, causes extensive interactions between the replication terminus and other genomic loci^31^. This observation is probably linked to the fact that the terminus is the major target region for Topo IV-mediated DNA decatenation^32^. Other Hi-C studies have shown that inactivation of DNA gyrase causes a global decrease in short-range (20-200 kb) interactions and more local structural changes at highly transcribed regions^33, 34^. The local effect is associated with the role of DNA gyrase in removing positive DNA supercoils ahead of RNA polymerase. In the archaeon *S. islandicus*, cold shock decreases plasmid linking number and alters global chromosome conformation, although the involvement of topoisomerases is unclear^17, 35, 36^.

Inspired by the above-mentioned findings, we hypothesized that 3C-based methodologies could illuminate the *in vivo* functions of Rgy, including its genomic targets. Using this approach, here we elucidate a key role of Rgy for maintaining the structural integrity of AT-rich sequences.

## Results

### Loss of Rgy has a limited effect on the 3D genome at permissive temperature

We have recently developed a high-resolution 3C-seq protocol for the hyperthermophilic euryarchaeon *T. kodakarensis*^37^. To obtain clues for the *in vivo* function of Rgy, we decided to apply this method to a *T. kodakarensis* strain lacking Rgy. Cells were grown to mid-to-late log phase at 85°C, where wild-type cells grow optimally while Δ*rgy* cells exhibit a modest decrease in growth rate^13^. As reported previously^37^, the genomic contact map from the parental strain KU216^38^ displayed a butterfly-like contact pattern resulting from a genomic inversion between two homologous proviral regions, *TKV2* and *TKV3*^39^ (Fig. 1a and Supplementary Fig. 1a). The KU216 genome also formed two boundaries (Fig. 1a), which have been named boundary 1 and boundary 2 respectively in our previous study^37^. The Δ*rgy* strain showed a similar but less intensive butterfly pattern (Fig. 1b and Supplementary Fig. 1b), probably because *T. kodakarensis* is polyploid and can possess both chromosome copies with and without the inversion at a variable ratio^37^. Differential contact analysis and contact probability plots revealed that the Δ*rgy* mutation resulted in a modest global decrease in short-range interactions up to 100 kb and a complementary increase in longer-range interactions (Fig. 1c, d). The Δ*rgy* mutant displayed similar levels of contact insulation across the two boundaries, although interestingly the mutant formed slightly intensified stripes at boundary 2 (Fig. 1e, f). Taken together, these results suggest that Rgy has a limited impact on genome architecture at 85°C.

**Fig. 1.**
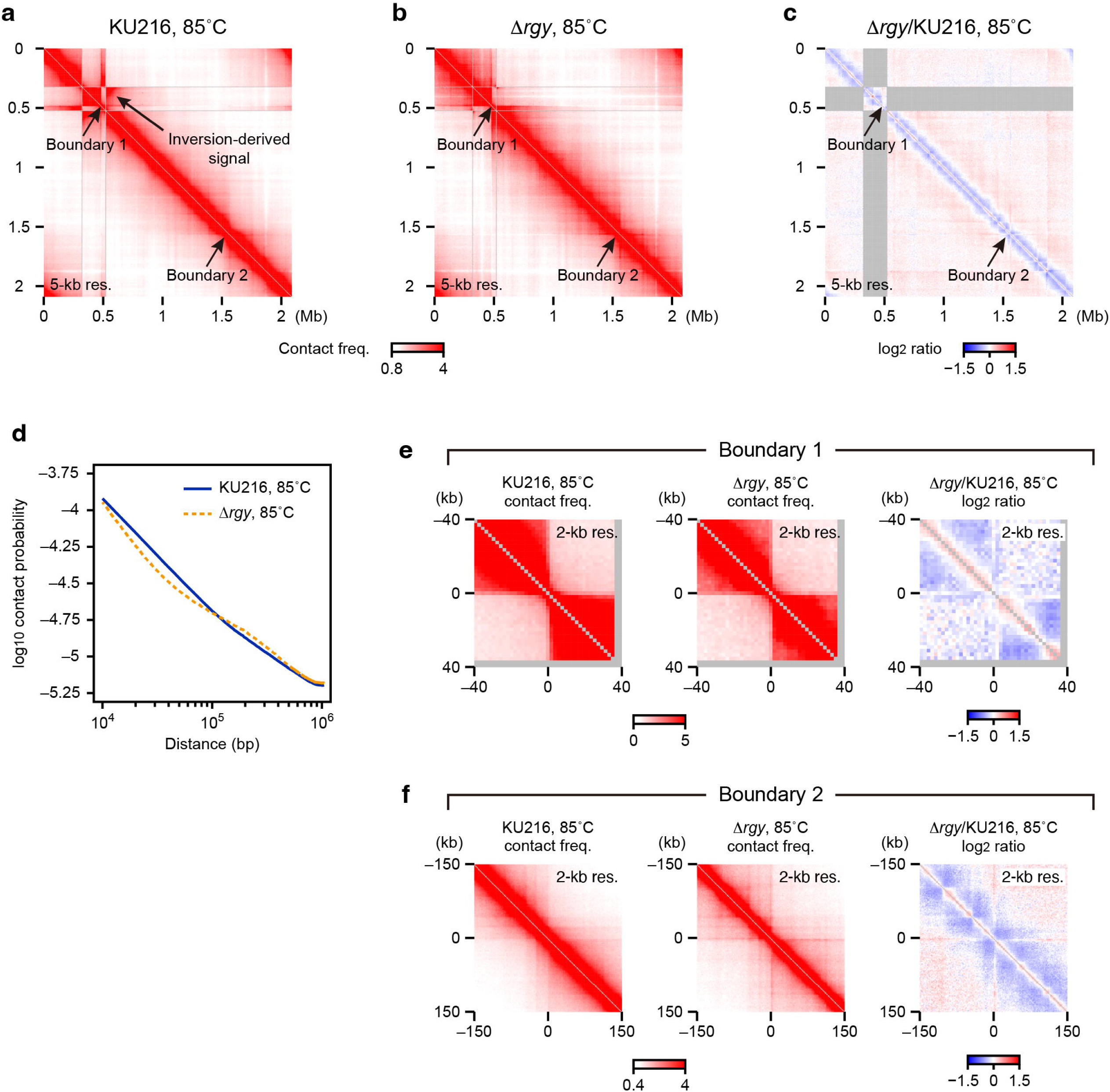
Loss of Rgy modestly decreases short-range interactions, while it has a very minor impact on domain insulation. **a, b** 3C-seq contact maps at 5-kb resolution. *T. kodakarensis* KU216 (**a**) and Δ*rgy* (**b**) cells were grown at 85°C. Two visible boundaries (boundary 1 and boundary 2) and an inversion-derived signal are indicated by arrows. **c** Differential contact map visualizing log_2_ ratios of contact frequencies shown in **a** and **b**. The genomic contacts between the inverted region and the other loci (shaded in gray) were omitted from the analysis. **d** Relative contact frequency averaged over 5-kb bin pairs of the same genomic distances (“contact probability”) was plotted as a function of genomic distance. Blue solid line: KU216 cells grown at 85°C. Dashed orange line: Δ*rgy* cells grown at 85°C. **e** Magnified views of **a-c** focusing on boundary 1 are shown at 2-kb resolution. **f** Magnified views of **a-c** focusing on boundary 2 are shown at 2-kb resolution.

We recently found that the formation of boundaries 1 and 2 requires Smc-ScpAB, a member of the SMC complexes^37^. Using a common ring-shaped structure, SMC complexes hold two segments of DNA together to organize genomes^40, 41^. A growing number of evidence suggest that SMC complexes use this biochemical activity to extrude DNA loops progressively^42^. Intriguingly, recent studies suggest that the dynamics of this loop extrusion process are controlled by DNA supercoiling^43, 44, 45, 46^. To investigate whether Smc-ScpAB activity is also modulated by Rgy, we performed ChIP-seq for the Smc subunit of the complex. At 85°C, the Smc distributions in KU216 and the Δ*rgy* strain were comparable at the genome-wide level and at boundaries 1 and 2 (Supplementary Fig. 2a-c). The identified Smc-binding sites were largely shared between the two strains, and these common binding sites displayed comparable Smc occupancy (Supplementary Fig. 2d, e). Thus, Rgy appears to play a minor role in regulating Smc-ScpAB. However, we cannot rule out the possibility that our ChIP-seq analysis failed to detect a global occupancy change due to the lack of a spike-in control.

### Loss of Rgy causes spatial gene clustering at non-permissive temperature

Given the minor effect of the Δ*rgy* mutation at 85°C, we decided to investigate how Rgy affects the 3D genome at a higher temperature. We grew cells to mid-to-late log phase at 85°C and then shifted the growth temperature to 95°C, which is permissive for wild type but not for Δ*rgy* cells^13^. One hour after the treatment, cells were fixed and used for 3C-seq. In response to the heat shock, KU216 and the Δ*rgy* strain displayed weakened contact insulation at boundaries 1 and 2 (Supplementary Fig. 3a-d). Their genomes also formed plaid-like contact patterns resulting from regional enrichment and depletion of long-range interactions (Fig. 2). The plaid pattern was more accentuated in the Δ*rgy* mutant due to a global increase in long-range contacts exceeding 200 kb compared to those in KU216 (Fig. 2b, d, and Supplementary Fig. 3e, f). The most striking observation was that, in the heat-shocked Δ*rgy* strain, several discrete loci were associated with one another extremely strongly, forming a dot grid in the plaid pattern (Fig. 2b, d). These results suggest that Rgy is required to suppress the clustering of specific genomic regions under heat stress.

**Fig. 2.**
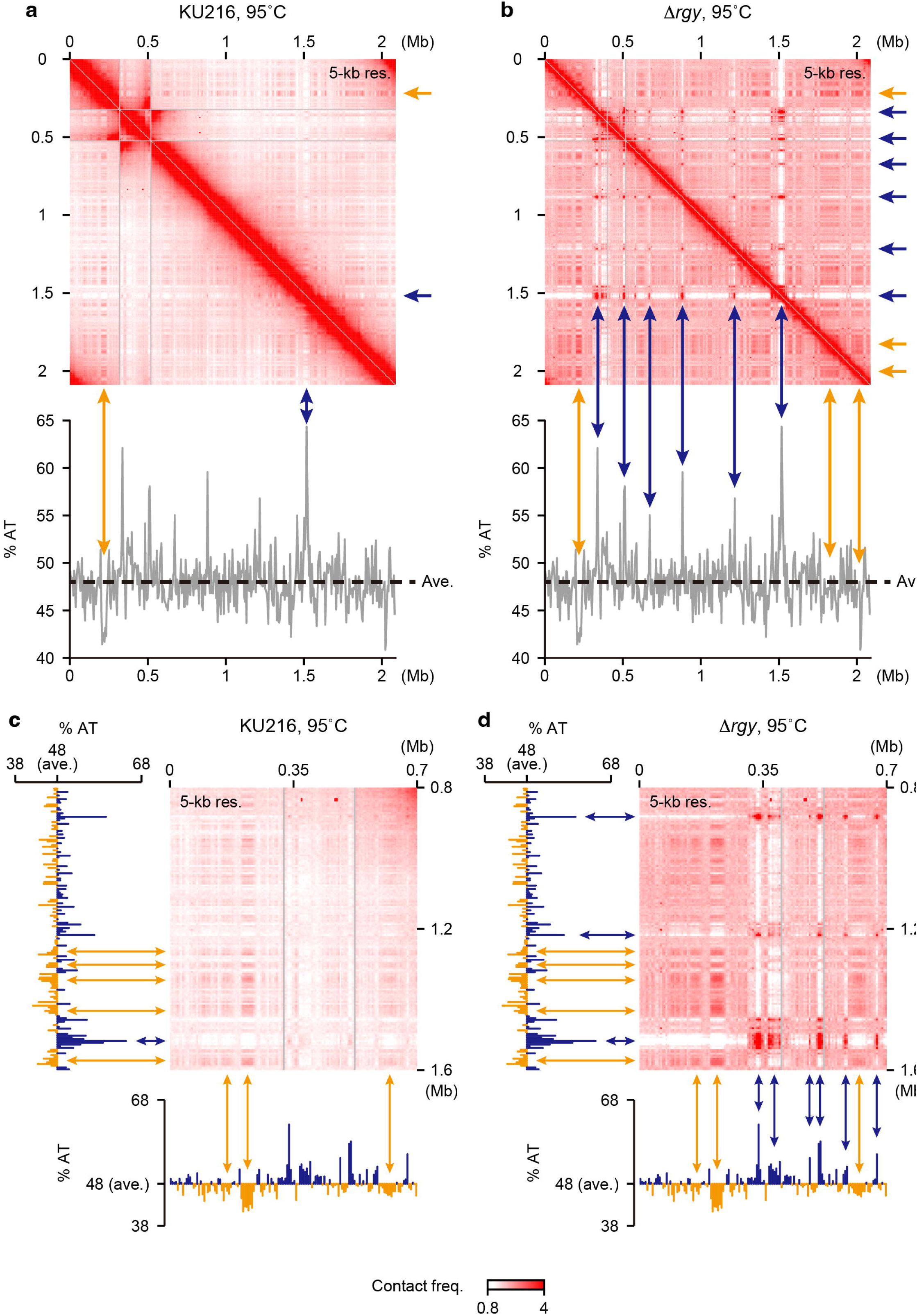
Δ*rgy* cells dramatically alter the genome conformation upon heat shock in a manner associated with genomic GC/AT content. **a, b** Top panels: 3C-seq contact maps at 5-kb resolution. KU216 (**a**) and Δ*rgy* (**b**) cells were exposed to 95°C for 1 h. Bottom panels: AT content was plotted over 5-kb bins. The average AT content of the genome is shown as dashed horizontal lines. Representative AT-rich and GC-rich regions forming stripes in either contact map are indicated by blue and orange arrows, respectively. **c, d** Magnified views of the contact maps in **a** and **b** are shown at 5-kb resolution, with AT-content tracks on the left and at the bottom at 5-kb resolution. Representative AT- and GC-rich regions forming stripes in either contact map are indicated by blue and orange arrows, respectively.

To investigate whether the gene clustering in the Δ*rgy* strain is indirectly caused by heat-induced cell death, we generated a *T. kodakarensis* strain lacking CpkB, a group II chaperonin critical for the viability of *T. kodakarensis* under heat stress^47^. At 85°C, the growth rate of the Δ*cpkB* mutant was comparable to that of the parental strain KU216, whereas at 95°C the former was significantly slower than the latter (Supplementary Fig. 4a, b). However, this heat sensitivity was milder than that previously reported for an isogenic Δ*cpkB* strain, possibly due to differences in the growth medium^47^. We next performed 3C-seq using our Δ*cpkB* mutant before and after a one-hour shift from 85°C to 95°C. At 85°C, the genomic contact matrices of KU216 and the Δ*cpkB* strain were largely similar to each other (Fig. 1a and Supplementary Fig. 4c). Upon the shift to 95°C, the contact matrix of the Δ*cpkB* srain exhibited a much more blurred plaid pattern than that of KU216 cells (Fig. 2a and Supplementary Fig. 4d). Importantly, the Δ*cpkB* strain did not form a dot-grid pattern of contacts at 95°C (Supplementary Fig. 4d). Thus, reduced viability itself is not the cause of the gene clustering phenotype under heat stress.

### Clustering occurs through genomic regions with high AT content

The mesophilic euryarchaeon *Haloferax volcanii* exhibits a weak plaid pattern of genomic contacts that correlates with GC/AT content^48^. This prompted us to investigate whether the genomic GC/AT content is associated with the heat-responsive gene clustering in *T. kodakarensis*. We found that, under the heat shock, GC-rich regions in both KU216 and the Δ*rgy* strain formed stripes through preferential self-interaction while excluding AT-rich loci (Fig. 2a, c). The heat-stressed Δ*rgy* strain was additionally characterized by discernible overlaps between AT-rich regions and dot-forming loci (Fig. 2b, d). To verify the spatial clustering of AT-rich sequences, we reordered 5-kb genomic bins by their AT content and visualized their contact frequencies relative to those expected from genomic distances. Generated distance-normalized contact matrices confirmed that, in the absence of heat stress, AT-rich regions had no contact bias in both KU216 and the Δ*rgy* strain (Fig. 3a, b). In contrast, we observed weak and much stronger clustering of the top several AT rich regions for KU216 and the Δ*rgy* strain, respectively (Fig. 3c, d). These results support the notion that loss of Rgy promotes the clustering of AT-rich loci under heat stress.

**Fig. 3.**
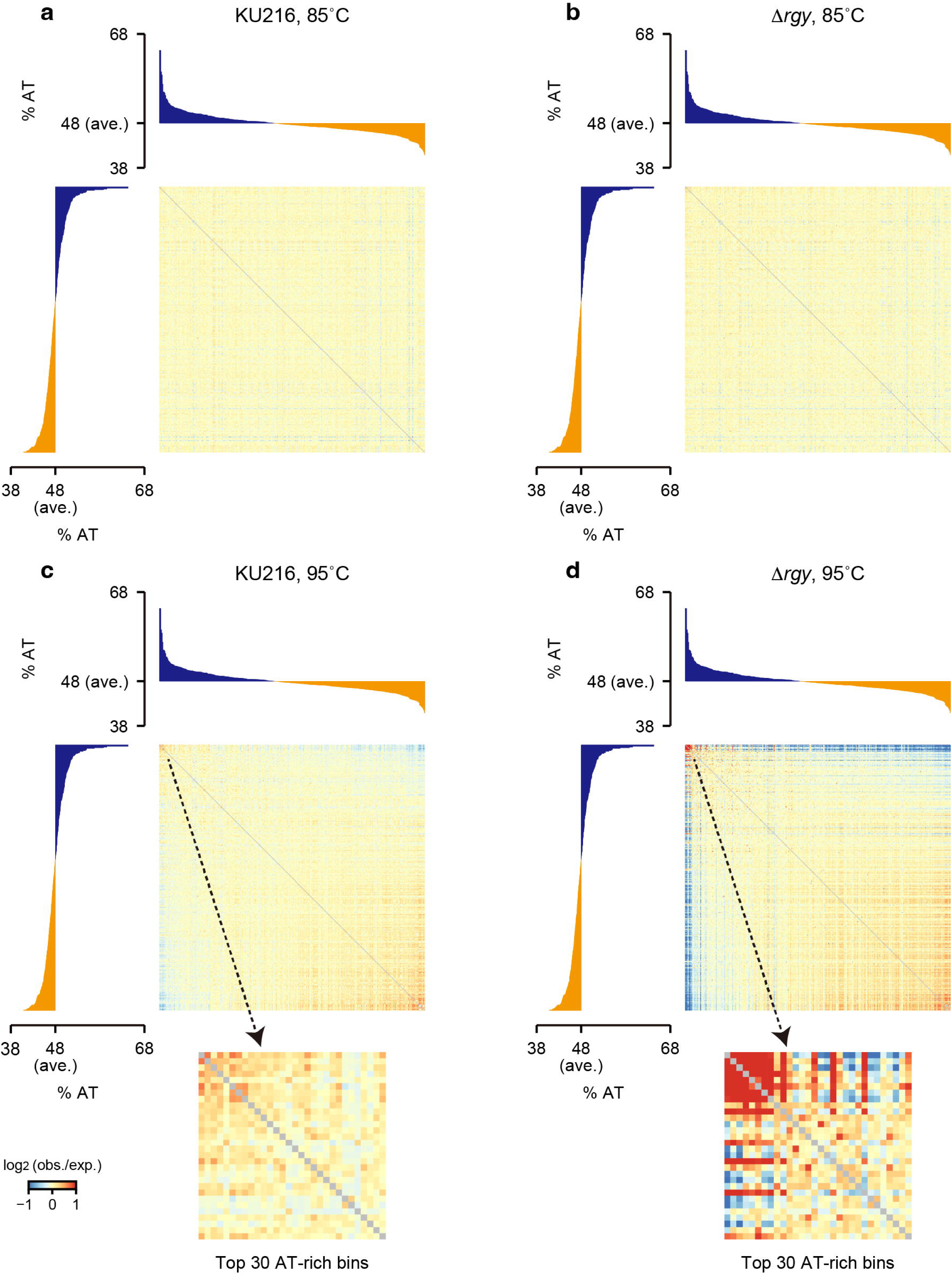
Loss of Rgy results in spatial clustering of AT-rich regions under heat stress. Distance-normalized contact matrices were generated at 5-kb resolution. KU216 and Δ*rgy* cells were grown at 85°C (**a** and **b**, respectively) and treated at 95°C for 1 h (**c** and **d**, respectively). Genomic bins were sorted by their AT content. For each matrix, a magnified view focusing on the top 30 AT-rich bins is also shown. AT-content tracks are shown on the left and on top of each matrix.

### The topoisomerase activity of Rgy is required to suppress the heat-induced gene clustering

Rgy can recognize nicked DNA and prevent DNA double-strand break (DSB) independently of its topoisomerase activity *in vitro*^21^. Given the potential non-enzymatic function of Rgy, we set out to determine whether its topoisomerase activity is required to suppress the clustering of AT-rich genomic regions. To this end, we generated a *T. kodakarensis* strain in which the catalytic Y963 residue of Rgy was substituted with phenylalanine. This mutation is hereafter denoted as *rgy^Y^*^963^*^F^*. The same mutation has been shown to abolish the *in vitro* topoisomerase activity of Rgy from the hyperthermophilic bacterium *Thermotoga maritima*^49^. Western blotting showed that *rgy^Y^*^963^*^F^* resulted in an approximately twofold increase in Rgy abundance (Fig. 4a, b). Despite this increase, *rgy^Y^*^963^*^F^* impaired cell growth partially at 85°C and completely at 95°C, similar to the phenotype observed with *rgy* deletion (Fig. 4c), suggesting that the topoisomerase activity of Rgy is essential to confer thermotolerance to *T. kodakarensis*. Next, we collected *rgy^Y^*^963^*^F^* cells before and after the temperature shift from 85°C to 95°C and used them for 3C-seq analysis. As observed for the *rgy* deletion, *rgy^Y^*^963^*^F^* caused a global decrease in short-range interactions and a complementary genome-wide increase in long-range interactions at both 85°C and 95°C (Supplementary Fig. 5a-d). Also as in the case with the Δ*rgy* mutation, *rgy^Y^*^963^*^F^* did not impair the formation of boundaries 1 and 2 at 85°C (Supplementary Fig. 5e, f), nor did it dampen the heat-induced disruption of the two boundaries (Supplementary Fig. 5g, h). Most importantly, *rgy^Y^*^963^*^F^* cells mimicked the heat-induced clustering of AT-rich loci observed for Δ*rgy* cells (Fig. 4d, e). Taken together, the topoisomerase activity of Rgy is required to prevent AT-rich regions from heat-induced clustering.

**Fig. 4.**
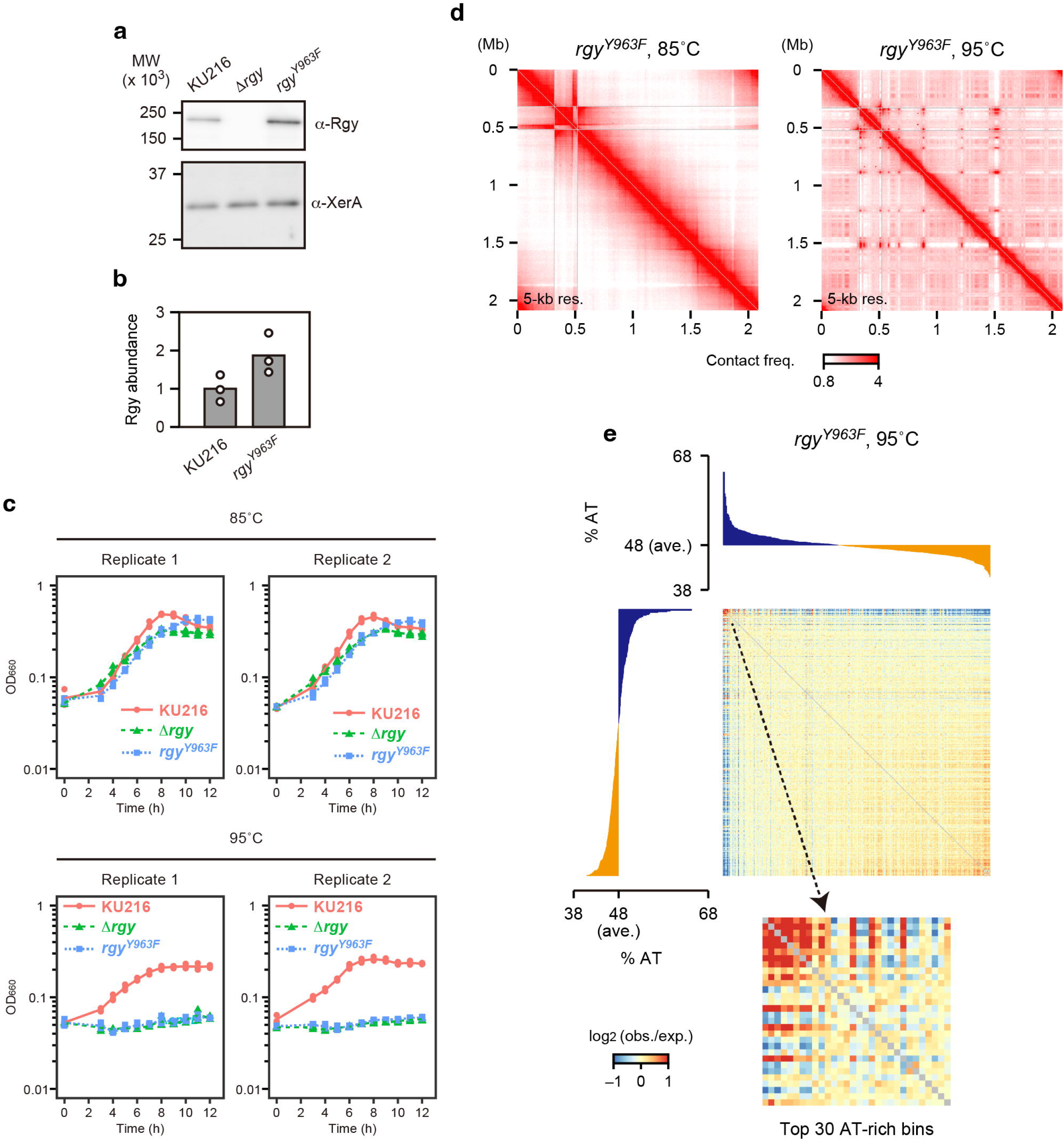
The DNA cleavage site of Rgy is required to prevent heat-induced clustering of AT-rich regions. **a** Western blot analysis of Rgy levels. XerA was used as a loading control. Molecular weight (MW) marker positions are indicated on the left. **b** Rgy abundance relative to that of XerA was quantified by western blotting. The quantification was performed for three biological replicates. **c** Growth curves of KU216 and derivative *rgy* mutants. For each replicate, a pre-culture was inoculated into three vials. These three cultures were used separately for OD_660_ measurement. The OD_660_ values were individually plotted (KU216: solid red circles, Δ*rgy*: solid green triangles, *rgy^Y^*^963^*^F^*: solid blue rectangles). A solid red line represents the mean OD_660_ values for KU216. Green and blue dashed lines represent the mean OD_660_ values for Δ*rgy* and *rgy^Y^*^963^*^F^*, respectively. **d** 3C-seq contact maps at 5-kb resolution. *rgy^Y^*^963^*^F^* cells were grown at 85°C (left panel) and treated at 95°C for 1 h (right panel). **e** A distance-normalized contact matrix from the *rgy^Y^*^963^*^F^* cells treated at 95°C for 1 h is shown as in Fig. 3, with AT-content tracks on the left and on top.

### AT-rich regions are prone to thermal denaturation in the absence of Rgy

It is reasonable to expect that the genomic regions with high AT contents are prone to thermal denaturation in the absence of Rgy-mediated DNA overwinding. All eukaryotes and the majority of archaea including *T. kodakarensis* possess Replication Protein A (RPA), a heterotrimeric ssDNA-binding protein complex^50, 51^. By taking advantage of the fact that RPA binding is a hallmark of ssDNA formation, we decided to map denatured DNA by performing ChIP-seq for Rpa41, a *T. kodakarensis* RPA subunit homologous to eukaryotic RPA70^52, 53^. Local enrichment of Rpa41 was not observed in KU216 cells regardless of the growth temperature, nor was it evident in Δ*rgy* cells at 85°C (Fig. 5a); however, the treatment of Δ*rgy* cells at 95°C induced Rpa41 binding to AT-rich genomic regions, which was increased sharply with AT content above ∼55% (Fig. 5a and b). Similar heat-induced binding was observed for Rpa32 and Rpa14, the other two subunits of *T. kodakarensis* RPA that are homologous to eukaryotic RPA32 and RPA14, respectively^53^ (Supplementary Fig. 6a-d). Rpa41, 32, and 14 shared most binding sites in heat-treated Δ*rgy* cells, corroborating that the three subunits act on the genome as a complex *in vivo* (Supplementary Fig.

**Fig. 5.**
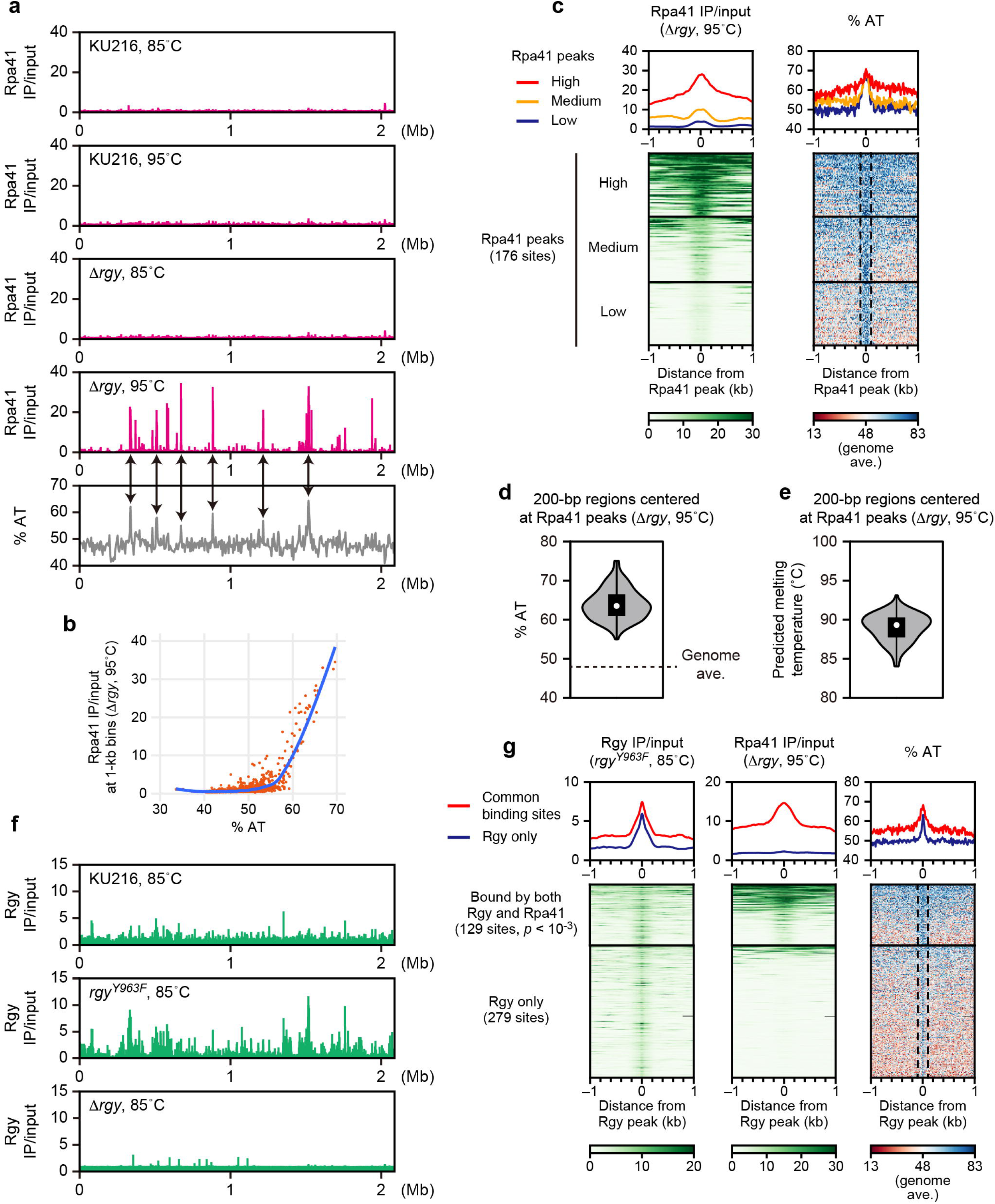
Loss of Rgy leads to hyper-accumulation of RPA at large AT-rich regions in response to heat shock. **a** Top four panels: ChIP-seq tracks of Rpa41. KU216 and Δ*rgy* cells were grown at 85°C (first and third from the top) and treated at 95°C for 1 h (second and fourth from the top). Enrichment of immunoprecipitated versus input DNA (IP/input) is shown at 1-kb resolution. Bottom: AT-content track at 5-kb resolution. Overlaps between high Rpa41 peaks and AT-rich regions are indicated by arrows. **b** AT content and Rpa41 IP/input values were plotted for each 1-kb bin. Blue line: a LOESS regression curve generated with a span of 0.2. **c** Left panels: average profiles (top) and heatmaps (bottom) of Rpa41 ChIP-seq signal were generated at 10-bp resolution. Δ*rgy* cells were treated at 95°C for 1 h. Rpa41 peaks were grouped based on the Rpa41 enrichment level. Right panels: average profiles (top) and heatmaps (bottom) of AT content were generated for the same regions as in the left panels. Dashed vertical lines highlight 200-bp regions centered at the Rpa41 peak summits. **d** Violin plot showing the AT content of 200-bp regions centered at the Rpa41 peak summits in the heat-treated Δ*rgy* cells. Dashed horizontal line: the average AT content of the genome. **e** Melting temperatures were predicted for the same 200-bp regions as in **d**. **f** Rgy ChIP-seq tracks in KU216, *rgy^Y^*^963^*^F^*, and Δ*rgy* cells grown at 85°C. The IP/input value is shown at 1-kb resolution. **g** Left panels: average profiles (top) and heatmaps (bottom) of Rgy ChIP-seq signal were generated at 10-bp resolution. *rgy^Y^*^963^*^F^* cells were grown at 85°C. Rgy peaks detected in the *rgy^Y^*^963^*^F^* cells were grouped based on their coincidence with Rpa41 peaks found in the Δ*rgy* cells treated at 95°C. Middle panels: average profiles (top) and heatmaps (bottom) of Rpa41 ChIP-seq signal were generated for the same regions as in the left panels using the data from the Δ*rgy* cells treated at 95°C. Right panels: average profiles (top) and heatmaps (bottom) of AT content were generated for the same regions as in the left panels.

6e). We focused on these common binding sites as the most probable RPA targets. These RPA-bound loci were AT-rich across 200 bp or larger areas centered at the Rpa41 peaks, displaying AT contents of 55 to 75% in the central 200-bp regions (Fig. 5c, d). The nearest-neighbor method^54^ predicted that most of these 200-bp RPA-bound regions have melting temperatures ranging from 85°C to 95°C (Fig. 5e), supporting the notion that they are denatured in the heat-treated Δ*rgy* cells. Intriguingly, strong Rpa41 peaks in the heat-stressed Δ*rgy* cells were distinguished by broadly high AT content rather than the AT richness of the peak summit (Fig. 5c). These peaks additionally exhibited broad enrichment of Rpa41 occupancy around the peaks. Taken together, loss of Rgy causes thermal denaturation of AT-rich regions at 95°C, especially genomic regions of ∼200-bp or larger with AT content of ∼55% or higher.

To verify that the melting-prone AT-rich regions are targeted by Rgy, we performed ChIP-seq with an antiserum raised against Rgy. Using Δ*rgy* cells grown at 85°C as a negative control, we identified 152 Rgy-binding sites in KU216 cells at the same temperature (Fig. 5f and Supplementary Fig. 7a). We also carried out ChIP-seq analysis of the *rgy^Y^*^963^*^F^* mutant grown at 85°C using the same antiserum. Intriguingly, Rgy^Y963F^ was associated with 138 of the 152 binding sites of wild-type Rgy with greater occupancy on average, while also localizing to 270 unique loci (Fig. 5f and Supplementary Fig. 7a, b). One plausible interpretation is that abolishment of the DNA cleavage step by *rgy^Y^*^963^*^F^* stabilizes the interaction between Rgy and the DNA substrate, which allowed for the identification of cryptic Rgy targets. We therefore used the Rgy^Y963F^ ChIP-seq data for downstream analyses, aiming to delineate a more comprehensive view of Rgy binding.

Rgy^Y963F^-bound loci were significantly overrepresented among the AT-rich, RPA-bound regions in the heat-stressed Δ*rgy* strain (Fig. 5g), consistent with the notion that Rgy targets AT-rich regions to prevent their thermal denaturation. We also identified a remarkable number of loci bound by Rgy^Y963F^ but not by Rpa41 (Fig. 5g). Of note, Rgy^Y963F^-bound regions, especially those not associated with Rpa41, formed sharper peaks of AT content than most Rpa41-bound regions did (Fig. 5c, g). This sequence difference can explain the different genomic distributions of Rgy and RPA.

### Clustering occurs through denatured AT-rich regions

To investigate whether thermal denaturation of AT-rich genomic regions is a prerequisite for their clustering, we examined the interactome of RPA-bound DNA fragments by HiChIP^55^ using the anti-Rpa41 antiserum. We detected no discernible point-to-point interactions in Δ*rgy* cells grown at 85°C (Supplementary Fig. 8). However, when Δ*rgy* cells were treated at 95°C, Rpa41-bound regions formed strong looping interactions (Fig. 6a). We determined the HiChIP loop positions using Chromosight^56^, observing that they recapitulated the dot-grid pattern in the 3C-seq contact map of Δ*rgy* cells at 95°C (Fig. 6b). This strongly supports the notion that AT-rich regions are clustered upon their denaturation.

**Fig. 6.**
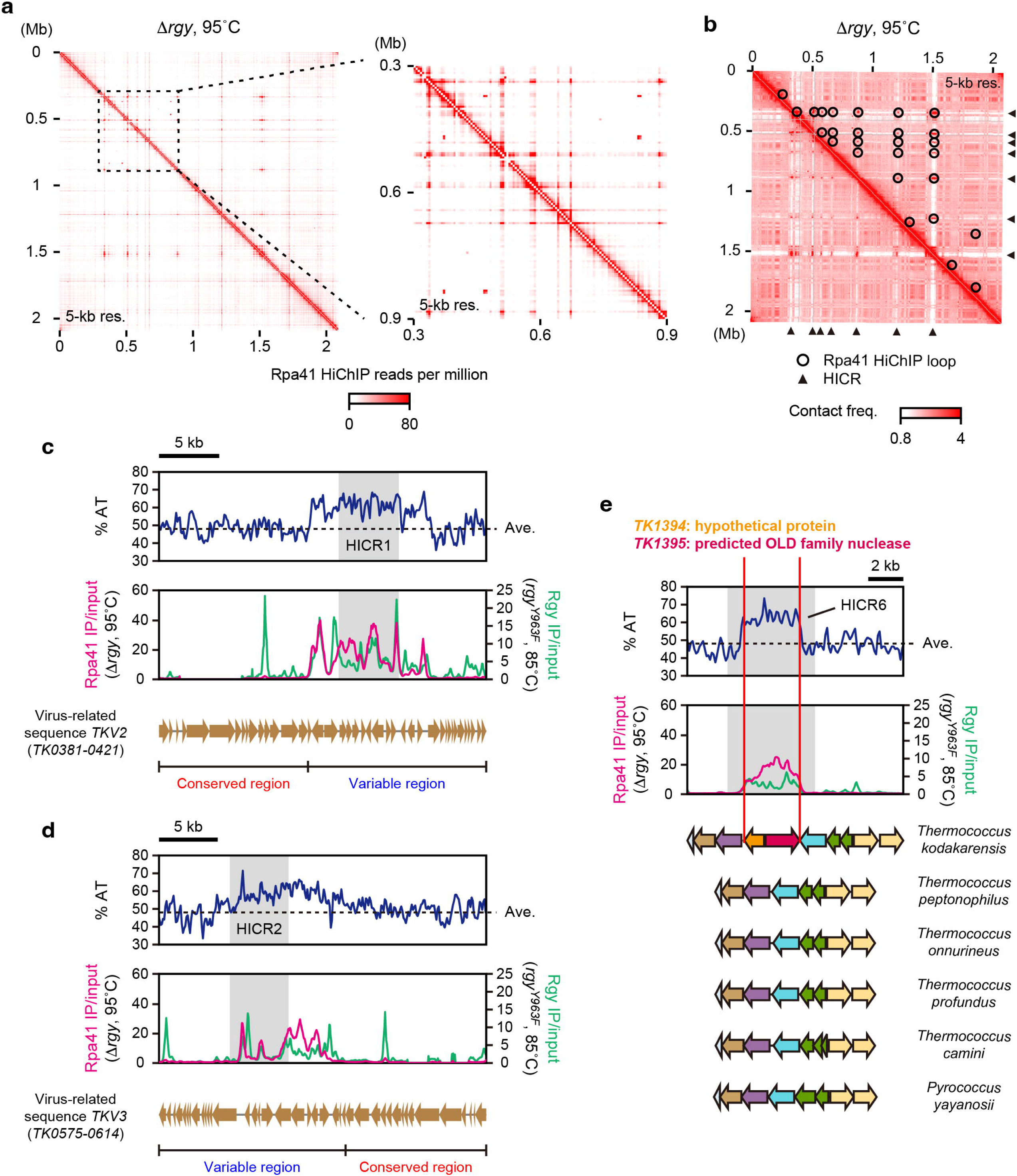
HICRs, AT-rich loci forming a spatial cluster in heat-stressed Δ*rgy* cells, are associated with signatures of horizontal gene transfer. **a** Left panel: Rpa41 HiChIP contact map 5-kb resolution. Δ*rgy* cells were treated at 95°C for 1 h. Right panel: a magnified view of the left panel, showing a representative dot-grid contact pattern indicative of gene clustering. **b** Significant looping interactions in **a** are shown by open circles on the same 3C-seq contact map as in Fig. 2b. Positions of the identified HICRs are shown by solid black triangles. **c, d** Genomic features of HICR1 (**c**) and HICR2 (**d**). Top panels: AT-content tracks were generated with a sliding window size of 200 bp and a step size of 100 bp. The average AT content of the genome is shown by dashed horizontal lines. Middle panels: an Rpa41 ChIP-seq track in Δ*rgy* cells treated at 95°C for 1 h (magenta) and an Rgy ChIP-seq track in *rgy^Y^*^963^*^F^* cells grown at 85°C (green) were generated at 50-bp resolution. In the top and middle panels, the locations of HICR1 and HICR2 are shaded in gray. Bottom panels: the annotated genes in *TKV2* (**c**) and *TKV3* (**d**) and their transcriptional directionality are indicated by arrows. Segments corresponding to conserved and variable regions in the pT26-2 family are indicated by horizontal lines with tick marks. **e** Genomic features of HICR6. AT-content and ChIP-seq tracks were generated as in **c** and **d**. In these tracks, the location of HICR6 is shaded in gray. Gene synteny was analyzed using Synttax^91^ and is indicated using colored arrows. Homologous genes are shown in the same color. Vertical red lines highlight the location of the non-conserved gene pair *TK1394* (orange arrow) and *TK1395* (red arrow).

By taking advantage of the high signal-to-noise ratio of the HiChIP data, we set out to determine the precise genomic positions of the clustered AT-rich loci. A spatial gene cluster can be seen as a clique of loop anchors that form loops with one another. We identified such a clique composed of seven HiChIP loop anchors, here referred to as HICRs (Heat-Induced Clustering Regions) (Fig. 6b). In the 3C-seq contact map of the heat-shocked Δ*rgy* cells, each of the seven HICRs strongly interacted with the other HICRs (Supplementary Fig. 9). All HICRs also overlapped with large AT-rich regions that were targeted by Rgy and became denatured in its absence (Fig. 6c-e, and Supplementary Fig. 10a-d). Thus, the HICRs recapitulated AT-rich loci forming the spatial cluster upon heat-induced denaturation.

### HICRs are associated with signatures of horizontal gene transfer

Genes that have been acquired via horizontal gene transfer (HGT) often exhibit atypical GC/AT content compared to the genome average^57^. Consistent with the extraordinary high AT content of the HICRs, many of them exhibit signatures of HGT. For instance, HICR1 and HICR2 (the seven HICRs are numbered sequentially according to the genomic coordinate) are located in the virus-related regions *TKV2* and *TKV3*, respectively^39^ (Fig. 6c, d). *TKV2*, *TKV3*, and some other virus-like sequences found in *Thermococcales* and *Methanococcales* share homology with the pT26-2 plasmid from *Thermococcus* sp. 26-2^58^. These homologous sequences are collectively called the pT26-2 family. Each member of this family can be divided into two DNA segments; one is highly conserved and the other is variable among the family. HICR1 and HICR2 are found in the variable regions of *TKV2* and *TKV3*, respectively. In another example, a large portion of HICR6 comes from the two adjacent genes *TK1394-1395* (Fig. 6e). This gene pair has broken a synteny block shared in members of *Thermococcales*, suggesting that the two genes have been integrated into the conserved gene cluster via HGT. Of note, *TK1395* exhibits homology with the OLD nuclease family, which constitutes diverse defense systems^59^. Current evidence suggests that many defense genes have been spread through HGT across prokaryotic lineages^60^, consistent with the potential horizontal acquisition of *TK1394-1395*. We also identified two other HICRs potentially related to defense systems; HICR4 contains *TK0773*, encoding another predicted member of the OLD nuclease family, and HICR5 overlaps with *TK1009-1010*, encoding a putative McrBC restriction-modification system^61^ (Supplementary Fig. 10b, c). Finally, IslandViewer 4^62^ revealed that HICR7 is located in a genomic island, a gene cluster with a probable horizontal origin (Supplementary Fig. 10d).

### RPA is required for efficient HICR clustering and thermotolerance in the absence of Rgy

A recent study has shown that eukaryotic RPA sequesters ssDNA by assembling into condensates^63^. This prompted us to investigate whether *T. kodakarensis* RPA plays a role for the spatial organization of HICRs. Prior work failed to delete the entire *rpa41-14-32* operon in *Thermococcus barophilus*, suggesting that the ssDNA-binding function of archaeal RPA is essential for viability^64^. Of note, *in vitro* studies demonstrated that Rpa14 is not required for ssDNA binding of archaeal RPA^52, 53^. We therefore sought to delete the potentially non-essential subunit Rpa14 and explore whether this deletion could perturb the HICR clustering.

A Δ*rpa14* mutant of *T. kodakarensis* was successfully obtained using KU216 as a parental strain. Although we reasoned that Rpa14 would be the most likely nonessential subunit of archaeal RPA, this finding was nonetheless surprising given that all RPA subunits are indispensable for cell viability in the eukaryote *Saccharomyces cerevisiae*^65^. Moreover, it was also possible to construct a Δ*rgy* Δ*rpa14* double mutant. Growth of both Δ*rpa14* and Δ*rgy* Δ*rpa14* mutants was moderately impaired at 85°C (Fig. 7a). 3C-seq analysis on these strains revealed that the Δ*rpa14* mutation caused a global decrease in short-range interactions up to around 100 kb and a complementary increase in longer-range interactions, regardless of growth temperature and the presence/absence of Rgy (Supplementary Fig. 11). The Δ*rgy* Δ*rpa14* strain formed the HICR cluster at 95°C, suggesting that Rpa14 is not essential for this genome reorganization (Fig. 7b). However, some of the heat-induced HICR-HICR contacts were modestly inhibited by the Δ*rpa14* mutation specifically in the Δ*rgy* background (Fig. 7c, d), pointing to a role of RPA in efficient HICR clustering.

**Fig. 7.**
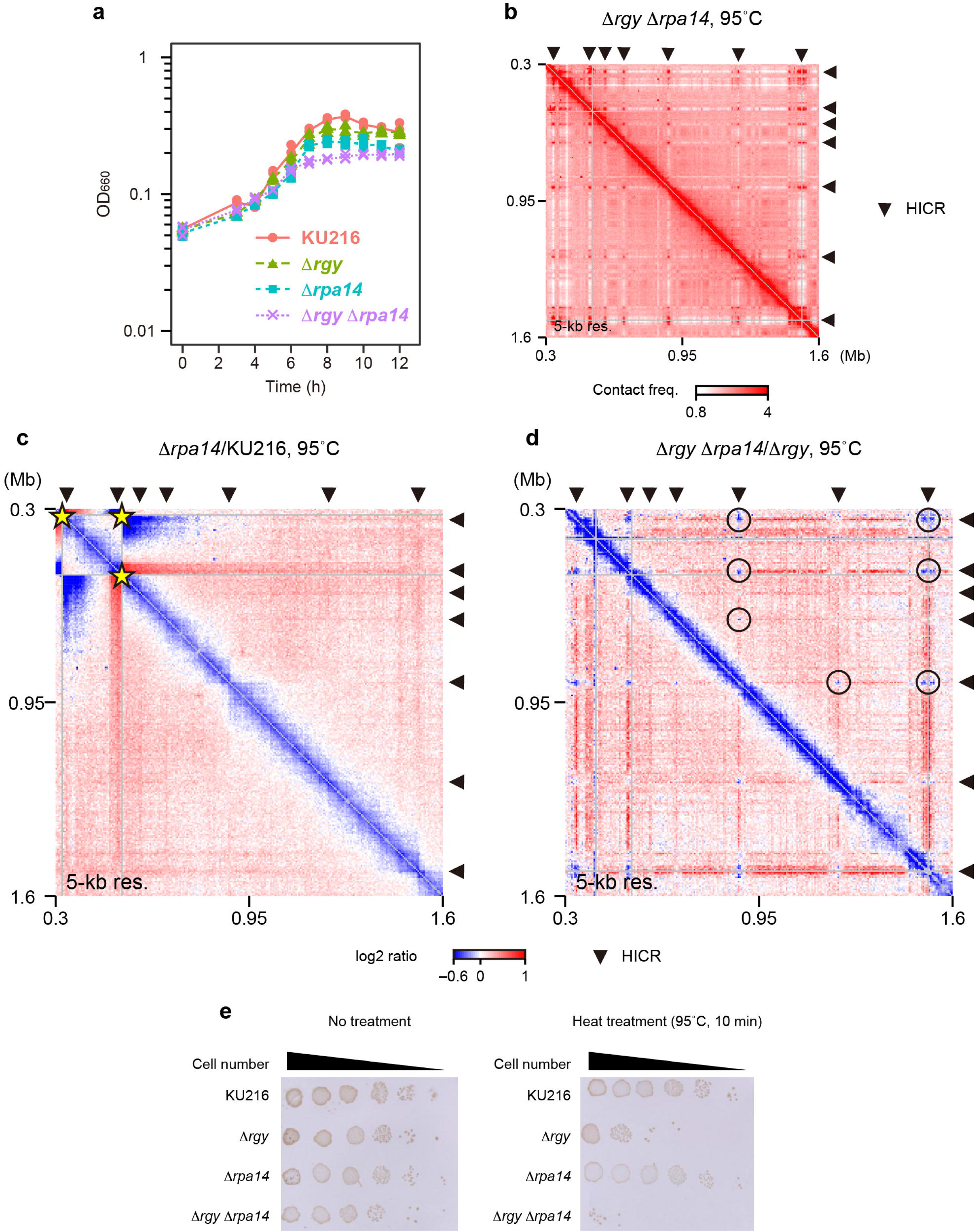
Loss of Rpa14 attenuates HICR clustering and exacerbates the heat sensitivity of Δ*rgy* cells. **a** Growth curves of KU216 and derivative mutants grown at 85°C. The data were collected from a single biological replicate, in which OD_660_ was measured as in Fig. 4c. The OD_660_ values were individually plotted (KU216: solid red circles, Δ*rgy*: solid green triangles, Δ*rpa14*: solid cyan rectangles, Δ*rgy* Δ*rpa14*: purple crosses). Solid and dashed lines represent the mean OD_660_ values for KU216 and the mutants, respectively (KU216: solid red line, Δ*rgy*: dashed green line, Δ*rpa14*: dashed cyan line, Δ*rgy* Δ*rpa14*: dashed purple line). **b** 3C-seq contact map showing interactions among the seven HICRs (solid black triangles) at 5-kb resolution. Δ*rgy* Δ*rpa14* cells were treated at 95°C for 1 h. **c, d** Differential contact maps visualizing log_2_ ratios of contact frequencies between indicated strains. The locations of the seven HICRs are shown by solid black triangles. In **c**, pronounced contact changes resulting from a genomic inversion are indicated by yellow stars. In **d**, markedly reduced HICR-HICR interactions are indicated by open circles. **e** Spotting assay was performed to examine the cell viability before (left panel) and after the treatment at 95°C for 10 min (right panel).

To examine whether the RPA-mediated coating or sequestration of denatured DNA regions contributes to the thermotolerance of *T. kodakarensis*, we compared the viability of KU216, Δ*rgy*, Δ*rpa14*, and Δ*rgy* Δ*rpa14* cells after heat stress. As cells lacking Rgy are expected to lose viability rapidly upon heat shock regardless of RPA, we exposed cells to 95°C for a short duration of 10 min. Whereas Δ*rgy* cells showed a pronounced growth defect in response to the heat shock, Δ*rpa14* cells did not markedly lose viability under the same conditions. However, the Δ*rpa14* mutation caused a synthetic growth defect in combination with the Δ*rgy* mutation upon the short heat treatment (Fig. 7e). These results suggest that RPA plays a key role in decelerating heat-induced cell death in the absence of Rgy.

## Discussion

Hyperthermophiles thrive in extremely hot habitats that can harm genome integrity^66, 67, 68, 69^. This extraordinary ability is thought to be conferred by several factors such as the DNA-stabilizing agent polyamine and abundantly expressed nucleoid-associated proteins^25, 70, 71^. Among them, Rgy has been hypothesized to play a particularly crucial role by generating positive DNA supercoils resistant to thermal denaturation, though without direct *in vivo* evidence^3, 10^. In this study, we have demonstrated that the topoisomerase function of Rgy is indeed required to prevent thermal denaturation of genomic DNA in the hyperthermophilic archaeon *T. kodakarensis* (Fig. 8).

**Fig. 8.**
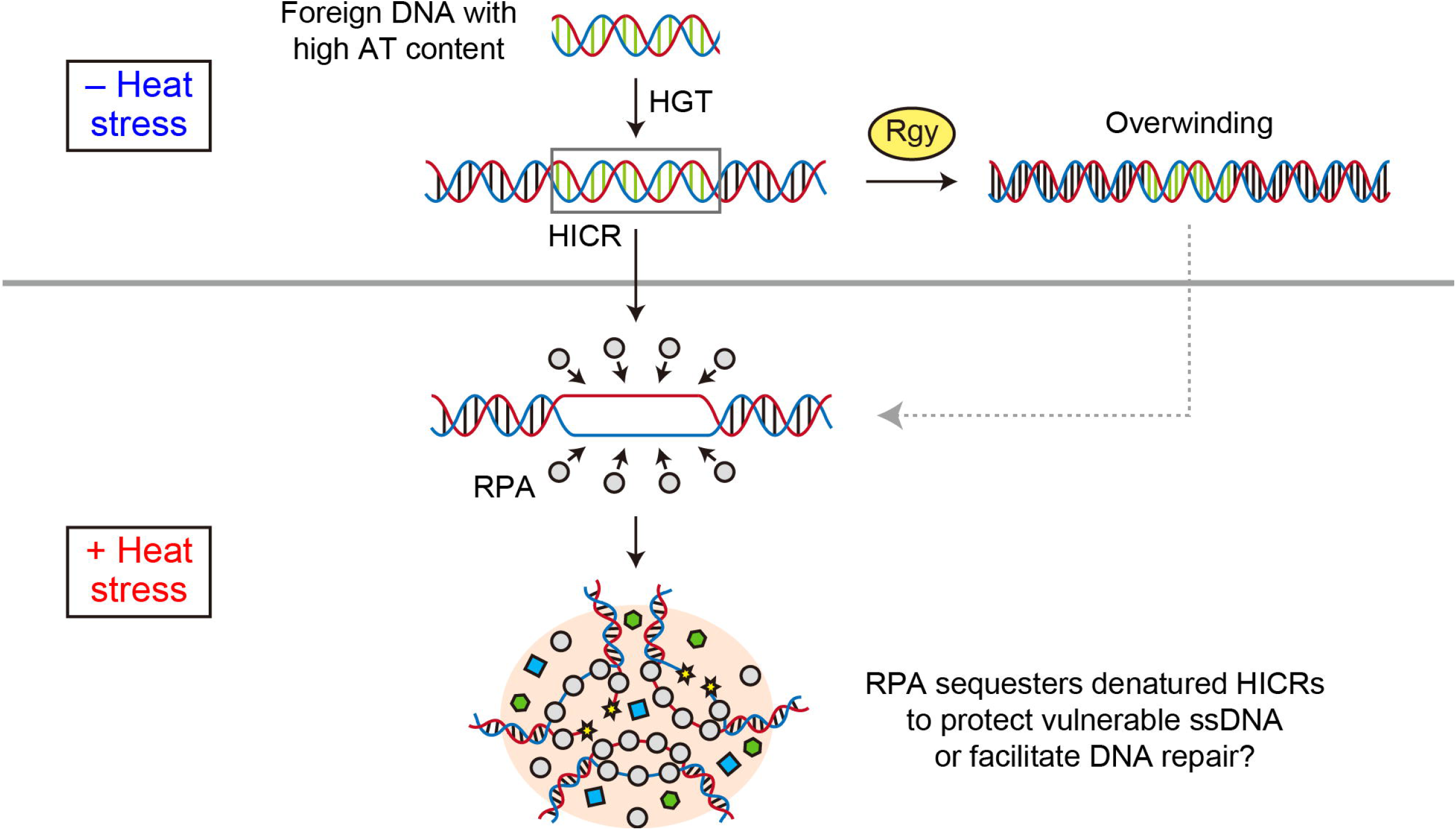
Model for the antagonistic interplay between Rgy-RPA axis and HGT in *T. kodakarensis*. Integration of foreign AT-rich sequences into the *T. kodakarensis* genome can compromise its stability under heat stress. In wild-type cells, this detrimental effect is counteracted by Rgy-mediated overwinding of AT-rich regions. When the function of Rgy is impaired, RPA sequesters denatured HICRs for their mechanical protection or repair.

Of note, our results uncovered that the Rgy-mediated suppression of DNA melting is highly localized to genomic regions with high AT content. This suggests that the impact of GC/AT content on genome thermal stability may need to be evaluated at a local scale, in contrast to earlier genomics studies showing that genome-wide GC content is at best only weakly correlated with optimal growth temperature^72, 73^. Consistent with the preferential action of Rgy on AT-rich sequences, a previous biochemical study demonstrated that Rgy-mediated positive supercoiling of plasmid DNA can be facilitated by a 38-bp insert of alternating AT dinucleotides. This work, which adopted an incubation temperature of 80°C, further showed that an artificial ssDNA bubble of similar size can also promote the positive supercoiling reaction^74^. Thus, Rgy likely recognizes denatured states of AT-rich regions, which can be as small as 38 nucleotides, rather than the sequences themselves. In contrast, RPA appears to require ∼200-nucleotide or larger ssDNA regions for stable binding, leading to its partially overlapping but different localization compared to that of Rgy (Fig. 5c, g).

Intriguingly, our data present a new example of how global genome organization is associated with DNA sequence features in euryarchaea. As observed in heat-stressed *T. kodakarensis* cells, the mesophilic halophilic euryarchaeon *H. volcanii* exhibits a plaid-like pattern of genomic contacts associated with GC/AT content. Our previous study also provided evidence that the nucleoid-associated protein TrmBL2 and specific AT-rich sequences known as A-tracts act as barriers to Smc-ScpAB in *T. kodakarensis*, thereby establishing a chromosomal domain boundary^37^. It is currently unknown whether these sequence-associated features of genome folding are mechanistically linked.

In recent work by Villain et al. (a preprint will be available shortly in *bioRxiv*), the authors investigated how Rgy controls the positive DNA supercoils in *T. kodakarensis* by trimethylpsoralen sequencing (TMP-seq), a DNA sequencing method that maps negatively supercoiled regions. They demonstrated that loss of Rgy increases TMP binding at promoters and transcription termination sites. This supports the notion that the topoisomerase activity of Rgy generates DNA positive supercoils, or at least reduces DNA negative supercoils, to stabilize AT-rich and histone-depleted genomic regions—such as promoters—against thermal denaturation. Prior work developed an *in vivo* mapping method for positive supercoils using GapR, a mesophilic bacterial protein that preferentially binds overtwisted DNA^75^. Future development of thermostable GapR through protein engineering may enable more direct investigation into whether and how Rgy introduces positive DNA supercoiling.

One of the most striking findings of this study is the identification of the HICRs, which not only become denatured but also spatially clustered in heat-stressed Δ*rgy* cells. Our data provide evidence that RPA extensively coats the HICRs and promotes their clustering. The HICR cluster may serve as a physical barrier that slows down the thermal degradation of vulnerable ssDNA, consistent with our finding that Δ*rgy* cells rapidly lose viability upon heat stress without Rpa14. It is also plausible that extensively denatured HICRs are brought together after breakage to facilitate DNA repair, reminiscent of the coalescence of DSBs into “repair centers” during eukaryotic DNA damage responses^76, 77^. In either case, we surmise that the higher-order assembly of HICR-RPA complexes presents a compensatory mechanism that safeguards the genome integrity of hyperthermophiles in case the Rgy function is compromised (Fig. 8).

How could RPA hold together the HICRs? Structural analysis has shown that the archaeal RPA can assemble into a tetrameric supercomplex in which two dimers of the Rpa41-32-14 trimer engage with each other via contacts between their Rpa14 subunits^53^. Although the tetrameric structure appears to be incompatible with ssDNA binding, it may indirectly bridge denatured HICRs by interacting with another HICR-binding protein. We also note that the HICR-RPA supercomplex is reminiscent of liquid droplets formed by human RPA and ssDNA^63^. Human RPA drives the phase separation using an intrinsically disordered region (IDR) situated at the N terminus of RPA2.

Intriguingly, archaeal Rpa32, a homolog of human RPA2, also possesses an IDR, although it is located between the N-terminal OB fold and the C-terminal winged helix domain^53, 78^. It is thus possible that Rpa32 plays an additional role in organizing HICR into a cluster.

Horizontally acquired genes can burden the host but also provide adaptive traits. Without Rgy, HGT-derived HICRs can exacerbate their fitness costs due to their denaturation-prone AT-rich sequences. Conversely, one can speculate that Rgy counteracts these detrimental effects of HGT on the genomic thermostability, thereby enabling hyperthermophiles to retain AT-rich foreign genes with adaptive potential. Notably, deletions of the homologous virus-related sequences *TKV2* and *TKV3* impair cell growth to varying degrees^79^, suggesting that their variable regions harboring HICRs contribute to fitness.

## Methods

### Cultivation of *T. kodakarensis* strains

Unless otherwise stated, cells were cultivated as follows. Cells were inoculated into ASW-YT-m1-S^0^ medium^80^ and grown overnight at 85°C. The overnight pre-culture was used to inoculated fresh ASW-YT-m1-S^0^ medium at an initial OD_660_ of 0.015. The culture was grown at 85°C until mid-log phase (OD_660_ ≈ 0.2) and then shifted to 95°C for 1 h. The redox indicator resazurin was added to ASW-YT-m1-S^0^ to a concentration of 0.5 mg/L.

### Construction of *T. kodakarensis* strains

The primers used for the construction are listed in Supplementary Data 1. The uracil-auxotrophic strain KU216^38^ (Δ*pyrF*) was used as a parental strain for strain construction.

Gene deletion and point mutation introduction was performed using a pop-in/pop-out approach^38^. For deletion of *rgy* (*TK0470*), two ∼1-kb regions immediately upstream and downstream of the coding sequence were PCR amplified using genomic DNA of *T. kodakarensis* KU216 as a template. The two fragments were cloned together into the EcoRI site of the pUD3 plasmid^81^ using In-Fusion HD Cloning Plus reagents (Takara Clontech, 638910). The obtained plasmid was named pUD3-Δ*rgy*. The plasmids for *cpkB* (*TK2303*) and *rpa14* (*TK1960*) deletion, named pUD3-Δ*cpkB* and pUD3-Δ*rpa14* respectively, were constructed the same way as for pUD3-Δ*rgy*. To introduce the Y963F substitution into *rgy*, a ∼3-kb fragment containing the target site was PCR amplified using genomic DNA of *T. kodakarensis* KU216 as a template. The fragment was cloned into the EcoRI site of pUD3 using In-Fusion HD Cloning Plus reagents. The obtained plasmid was further used as a template for inverse PCR using a primer set containing the Y963F substitution. The PCR fragment was circularized using In-Fusion HD Cloning Plus reagents. The obtained plasmid was named pUD3-*rgyY963F*. The generated plasmids were used for transformation as described previously^37^.

### Growth measurement of *T. kodakarensis*

Growth measurement was carried out as described previously^37^.

### Spotting assay

*T. kodakarensis* culture was grown at 85°C until mid-log phase (OD_660_ ≈ 0.2) and then shifted to room temperature for approximately 30 min until the culture cooled down. The following procedures except for centrifugation were performed in an anaerobic chamber. The culture was diluted in 0.8 × ASW-m1^37^ to an OD_660_ of 0.15 for a total volume of 5 mL. Cells were harvested by centrifugation (10,000 × *g*, 5 min, 25°C) and resuspended in 750 μL of 0.8 × ASW-m1. 200 μL of the suspension was transferred to each of two 1.5-mL tubes containing 800 μL of ambient 0.8 × ASW-m1. One of the dilutions was subjected to heat stress by incubating for 10 min at 95°C with agitation at 600 rpm. The heated cells were immediately cooled down on ice for 5 min. The other dilution was left at room temperature without agitation. Each of the two samples was centrifuged (10,000 × *g*, 5 min, 25°C), resuspended in 100 μL of 0.8 × ASW-m1, and serially diluted ten-fold in 0.8 × ASW-m1. 2 μL of each dilution were spotted on a ASW-YT-m1 plate and anaerobically incubated at 85°C for 20 h. The cells on the plate were transferred to a PVDF membrane and imaged. Experiments were repeated three times independently.

## 3C-seq

3C-seq was carried out as described previously^37^.

### Protein purification and antibody production

The coding region of *TK0470* (*rgy*) without an intein-coding sequence was expressed using EasySelect™ Pichia Expression Kit (ThermoFisher, K174001) according to the manufacturer’s instructions. The cell lysate was suspended in a buffer A (50 mM Tris-HCl [pH 8.0], 0.1 mM EDTA) containing 0.5 M NaCl. After centrifugation, the supernatant was incubated at 80°C for 20 min, and the heat-resistant fraction was obtained by another centrifugation. Polyethyleneimine was added to precipitated nucleic acids in the heat-resistant fraction at a concentration of 0.15% (w/v). The supernatant was subjected to ammonium sulfate precipitation. Proteins in the solution was precipitated by 80% saturation of ammonium sulfate, and were resuspended in buffer A containing 1 M (NH_4_)_2_SO_4_, then applied to a HiTrap^TM^ Butyl HP column (Cytiva, 28411005). Hydrophobic interaction chromatography was performed with a 1–0 M (NH_4_)_2_SO_4_ gradient in buffer A. The Rgy-containing fraction was dialyzed against buffer A containing 0.1 M NaCl, and then applied to a HiTrap^TM^ Heparin HP column (Cytiva, 17040701), which was developed with a linear gradient of 0.1–1 M NaCl in buffer A. The eluted Rgy was subjected to size-exclusion chromatography using a Superdex^TM^ 200 16/600 column (Cytiva, 28989335) equilibrated with buffer A containing 0.3 M NaCl. The purified protein was provided as an antigen to obtain an antiserum (Eurofin Genomics).

### ChIP-seq

ChIP-seq was carried out as described previously^37^ using the antisera against Rpa41, Rpa32, and Rpa14 generated previously^52^ and the anti-Rgy antiserum raised as described above.

### HiChIP

#### Cell Fixation

1. *T. kodakarensis* cells were grown and crosslinked as in 3C-seq. The pellet was stored at –80°C until use.

#### Restriction digestion

Crosslinked cells were processed as in 3C-seq with the following modifications. For DNA digestion, 90 μL of the lysate was mixed with premixed buffer composed of 41 μL of 10 × NEBuffer 2 (New England Biolabs, B7002S), 150 μL of 10% v/v Triton X-100, and 209 μL of MilliQ water. This was further mixed with 10 μL of 25 U/μL MboI (New England Biolabs, R0147M) and incubated for 3 h at 37°C with agitation at 600 rpm. The tube was also inverted every 30 min for mixing. The reaction mixture was heated for 20 min at 65°C with agitation at 600 rpm to inactivate the restriction enzyme. 25 μL of the reaction mixture was stored for purification of un-ligated control DNA.

#### Proximity ligation

The insoluble fraction was collected by centrifugation (16,000 × *g*, 10 min, 25°C), resuspended in 500 μL of wash buffer (10 mM Tris-HCl [pH 8.0], 50 mM NaCl, 10 mM MgCl_2_), and centrifuged again (16,000 × *g*, 10 min, 25°C). The pellet was resuspended in 790 μL of 1.01 x T4 DNA Ligase Reaction Buffer (New England Biolabs) and mixed with 10 μL of 400 U/μL T4 DNA Ligase (New England Biolabs, B0202S) and incubated for 3 h at 16°C with agitation at 600 rpm. The reaction was also mixed every 30 min by inverting the tube. 40 μL of the reaction mixture was stored for purification of un-sonicated control DNA.

#### Sonication

The insoluble fraction was collected by centrifugation (16,000 × *g*, 10 min, 25°C), resuspended in 600 μL of TBS-TT (20 mM Tris-HCl [pH 8], 150 mM NaCl, 0.1% v/v Triton X-100, and 0.1% v/v Tween 20), and transferred to a milliTUBE 1 ml AFA Fiber (Covaris, 520135) for DNA fragmentation using M220 Focused-ultrasonicator (Covaris). The fragmentation was performed for 400 sec at 7°C with the peak power set to 75, the duty factor set to 26, and the cycles/burst set to 200. The lysate was transferred to an 1.5-mL tube and centrifuged (20,400 × *g*, 15 min, 4°C). 25 μL of the supernatant was stored for purification of sonicated control DNA.

#### Immunoprecipitation

500 μL of the lysate was mixed with 5 μL of antiserum and gently rotated overnight at 4°C. The sample was further mixed with 50 μL of Dynabeads Protein G (Thermo Fisher Scientific, 10004D) pre-washed with TBS-TT. The lysate-bead mixture was gently rotated for 1h at 4°C. The beads were washed three times with TBS-TT, once with TBS-TT containing 0.5 M NaCl, and once with TBS-TT containing 0.5% v/v Triton X-100 and 0.5% v/v Tween 20.

Protein-DNA complexes were eluted in 200 μL of elution buffer (20 mM Tris-HCl [pH 8.0], 10 mM EDTA [pH 8.0], and 0.5% w/v SDS) for 30 min at 65°C with agitation at 1,400 rpm. The eluted sample was mixed with 3 μL of 800 U/mL proteinase K (New England Biolabs, P8107) overnight at 37°C. The DNA was extracted with phenol:chloroform:isoamyl alcohol (Sigma Aldrich, 77618-100ML) two times and ethanol-precipitated together with 2 μL of 20 mg/mL glycogen (Thermo Fisher Scientific, R0561). The DNA was dissolved in 53 μL of 10 mM Tris-HCl (pH 8.0).

#### Library construction

The purified DNA was used for library construction with NEBNext Ultra II DNA Library Prep with Sample Purification Beads (New England Biolabs, E7103S). Reaction was performed according to the manufacturer’s instructions with size selection for a 300-400 bp insert.

### Illumina DNA sequencing and read mapping

The 3C-seq, ChIP-seq, and HiChIP libraries generated in this study were paired-end sequenced on either of the Illumina NovaSeq 6000 and NextSeq 550 platforms. The sequencing was performed by Single-Cell Genome Information Analysis Core at Kyoto University. Generated data were mapped to the reference genome of *T. kodakarensis* KOD1 (GCA_000009965.1).

### 3C-seq data analysis

#### Generation of contact matrices

Iteratively corrected contact matrices were generated as described previously^37^ using HiC-Pro (version 3.1.0)^82^. The obtained matrices were normalized so that the sum of interaction scores is equal to 1000 for each row and column, and these normalized values were shown as contact frequencies. Distance-normalized contact matrices were generated by calculating the log_2_ ratio of the observed contact frequency to the expected value (the genome-wide average contact frequency for the corresponding genomic distance) for each pair of bins.

Differential contact matrices were generated by calculating the log_2_ ratio of contact frequencies between the two samples. In most differential contact matrices, the genomic contacts between the bins overlapping the inversion (the genomic coordinates 320001–525000) and the other bins were omitted.

#### Insulation score

Insulation score was determined as described previously at 1-kb resolution^37^.

### ChIP-seq data analysis

#### Mapping

Reads were mapped using Bowtie 2 (version 2.4.4)^83^. Alignments with MAPQ < 30 were removed using SAMtools (version 1.12)^84^.

#### Generation of ChIP-seq tracks

BAM alignment files were processed using the bamCoverage function (version 3.5.4) of deepTools^85^ to calculate Reads Per Kilobase region per Million mapped reads (RPKM) for genomic bins of 50 bp and 1 kb. RPKM ratios of immunoprecipitated versus input DNA were plotted as IP/input.

#### Peak analysis

Peak calling was performed as described previously using MACS (version 3.0.1)^86^. To identify overlapping peaks between multiple datasets, overlaps of ±100-bp regions from the peak summits were examined using the venn function of intervene (version 0.6.5)^87^. Statistical significance of the overlaps was tested using the overlapPermuTest function of regioneR (version 1.30.0)^88^ with ntimes = 1000. The Rpa41 peaks that overlap both Rpa32 and Rpa14 peaks were regarded as bona fide Rpa41-bound regions for downstream analysis. The IP/injput ratio of an Smc peak was defined as (*C_IP_*/*C_input_*) × (*N_IP_*/*N_input_*), where *C* is the read coverage for the ±100-bp region from the peak summit and *N* is the total number of reads in the library. The subscripts *IP* and *input* represent immunoprecipitated and input DNA respectively.

#### Average profiles and heatmaps

Average profiles and heatmaps of ChIP-seq signals around regions of interest were generated as follows. First, IP/input tracks were generated using the bamCompare (version 3.5.6) function of deepTools with the following parameters: *-bs 10 --scaleFactorsMethod readCount --operation ratio*. Generated bigWig files were processed using the computeMatrix function of deepTools with the following parameters: *-b 1000 -a 1000 --referencePoint TSS*. Generated gz files were processed using the plotHeatmap (version 3.5.6) function of deepTools. A bigWig file containing the AT content of each 10-bp bin was processed in the same way.

### HiChIP data analysis

#### Generation of contact matrices

HiChIP contact matrices were generated at 5-kb resolution as described for the 3C-seq contact matrices but without iterative correction. The read counts in the matrix were normalized to one million mapped reads.

#### Identification of the HICRs

HICRs were defined as a group of loop anchors that become denatured and form loops with one another in heat-shocked Δ*rgy* cells. To identify such loci, loop positions in the Rpa41 HiChIP data were first determined using Chromosight (version 1.6.3)^56^. We then defined the strength of the loop between the *i*th and *j*th bins as the median of the read counts in the 3 × 3 submatrix centered at the *i*th row and *j*th column. The loops with a strength of 15 or greater were retained and visualized in Fig. 6b. Among the retained loop anchors, those that participated in multiple loops were defined as HICRs. This definition does not necessarily guarantee that the identified HICRs form loops with all the other HICRs. Indeed, among all the 21 possible HICR-HICR loops, we failed to detect those of the HICR1-2 and HICR2-7 pairs. However, HICR1 and HICR7 instead formed loops with bins proximal to HICR2, which are only 10-kb and 5-kb away from HICR2 respectively. We also note that these HICR pairs, together with their neighboring regions, made strong contacts in the 3C-seq data of heat-treated Δ*rgy* cells (Supplementary Fig. 9). We thus conclude that all the identified HICRs coalesce into a spatial cluster.

### AT content analysis

AT content was calculated for 10-bp bins using BEDtools (version 2.31.1)^89^ and for 200-bp and 5-kb bins using SeqKit (version 2.5.1)^90^. The AT-content track at 200-bp resolution was smoothed by setting the sliding window size to 100 bp. The AT content of Rpa41 peaks was calculated using SeqKit.

### Melting temperature prediction

DNA melting temperature was predicted for ±100-bp regions from the Rpa41 peak summits using the following formula based on the nearest-neighbor method^54^:

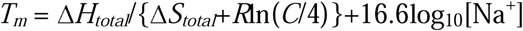

where *T_m_* is the melting temperature, Δ*H_total_* (kcal/mol) is the sum of the nearest-neighbor enthalpy changes for all base pair steps in the sequence, Δ*S_total_* (cal/mol·K) is the sum of the nearest-neighbor entropy changes for all base pair steps in the sequence, *R* is the gas constant (1.987 cal/mol·K), *C* is the concentration of the DNA sequence. In this study, *C* was set to 2 nM and [Na^+^] (mol/L) was set to 0.7. [Na^+^] was included in the formula to adjust for the effect of salt concentration on DNA melting.

### Western blotting

10 mL of cell culture were grown at 85°C until mid-log phase (OD_660_ ≈ 0.2) and transferred to a 50-mL tube containing 20 mL of pre-chilled 0.8 × ASW-m1. After cooling in iced water for 3 min with occasional agitation, the cells were centrifuged (10,000 × *g*, 4°C, 10 min). The pellet was resuspended in 1 mL of pre-chilled 0.8 × ASW-m1 and centrifuged (10,000 × *g*, 4°C, 5 min). The pellet was resuspended in 500 μL of lysis buffer (50 mM Tris-HCl [pH 8.0], 0.5 M NaCl, 0.1% v/v Triton X-100, and 0.1% v/v Tween-20) and transferred to a milliTUBE 1 mL AFA Fiber (Covaris, 520135) for sonication using M220 Focused-ultrasonicator (Covaris). The cell disruption was performed for 400 sec at 7°C with the peak power set to 75, the duty factor set to 26, and the cycles/burst set to 200. The lysate was transferred to an 1.5-mL tube and centrifuged (20,400 × *g*, 4°C, 15 min). The supernatant was diluted in lysis buffer to a protein concentration to 2.3 mg/mL. 395 μL of the diluted sample were mixed with 100 μL of 10% w/v SDS and 5 μL of 2-mercaptoethanol. The mixture was heated for 10 min at 95°C and stored at –20°C until use. The lysate containing 7.2 μg of proteins was used for western blotting. Blots were imaged using ImageQuant LAS-500 (Cytiva). The band intensity was quantified using ImageJ (version 1.52v).

### Data visualization and statistical test

Unless otherwise stated, data were visualized using R software (http://www.r-project.org/).

### Statistics and reproducibility

Statistical test was performed using R software. Reproducibility of 3C-seq was confirmed using five (KU216 cells grown at 85°C), four (KU216 cells treated at 95°C), or two (the others) biological replicates. Reproducibility of HiChIP was confirmed using two biological replicates.

Reproducibility of ChIP-seq was confirmed using two biological replicates. Reads from the biological replicates were pooled for high-throughput DNA sequencing analysis. Growth measurement was performed once or twice independently with similar results. Quantification of western blots was performed for three biological replicates.

## Data Availability

## Acknowledgements

We thank Single-Cell Genome Information Analysis Core (SignAC) at WPI-ASHBi at Kyoto University for Illumina DNA sequencing. We also thank Tamara Basta for sharing unpublished data.

K.Y. is supported by JST, the establishment of university fellowships towards the creation of science technology innovation (JPMJFS2123) and SPRING (JPMJSP2110). N.T. is funded by JST FOREST (JPMJFR224V), JSPS Grant-in-Aid for Early-Career Scientists (JP22K15083), and JSPS Grants-in-Aid for Transformative Research Areas (JP23H04281).

## Author contributions

K.Y. performed strain construction, growth measurement, 3C-seq, HiChIP, ChIP-seq, and western blotting. K.Y. and N.T. performed data analysis. T.Y. and Y.I. performed protein purification to raise the anti-Rgy antiserum for ChIP-seq. N.T. and H.A. supervised the project. N.T. and Y.I. acquired funding. K.Y. and N.T. wrote the manuscript. All co-authors critically read and edited the manuscript.

## Additional information

**Correspondence** and requests for materials should be addressed to Naomichi Takemata or Haruyuki Atomi.

