## Supplementary Figure for "Reverse gyrase and 3D genome architecture suppress hyperthermophile genome instability arising from horizontal gene transfer"

### Supplementary Figures

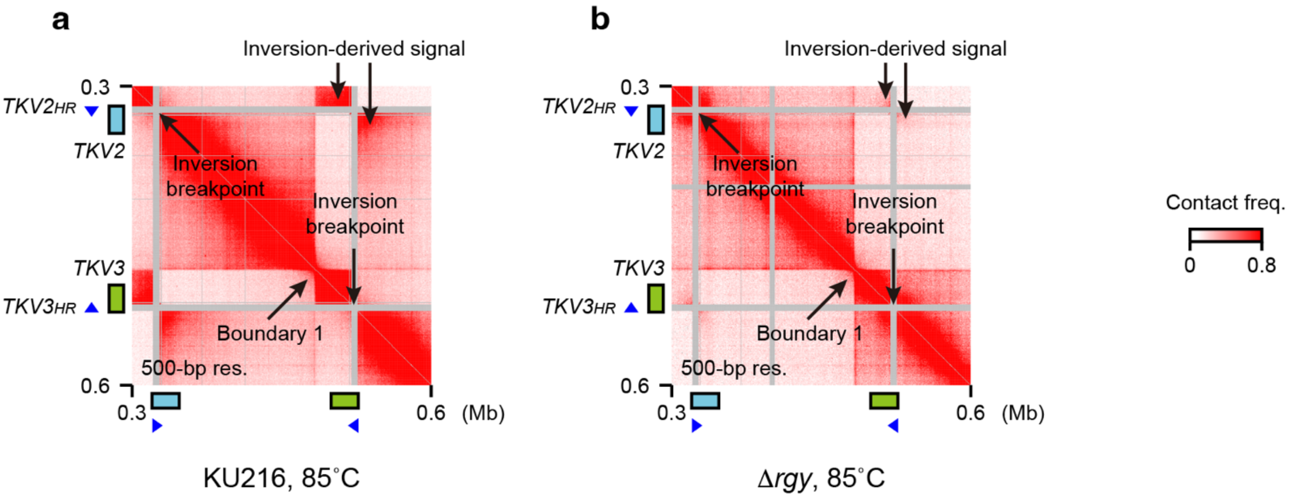


**Supplementary Figure 1 | Inversion-derived contacts in KU216 and Δ*rgy* cells**

Magnified views of Fig. 1a (**a**) and b (**b**) were generated at 500-bp resolution to show a butterfly pattern indicative of a genomic inversion. Two proviral regions (*TKV2* and *TKV3*) are indicated by cyan and green rectangles, respectively. Homologous sequences within *TKV2* and *TKV3* (*TKV2*$^{HR}$ and *TKV3*$^{HR}$, respectively) are indicated by blue triangles. Other features are highlighted by arrows.

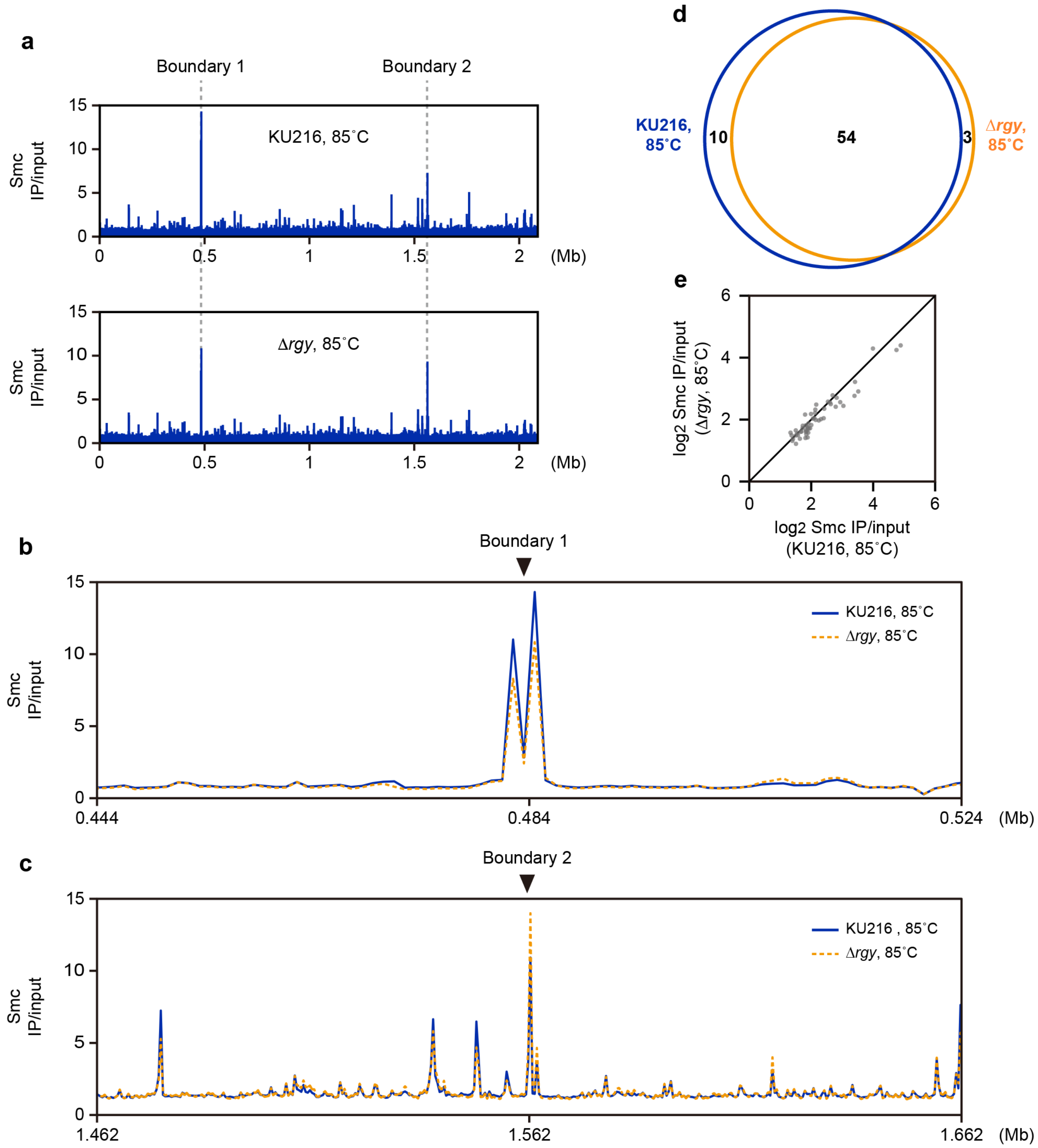


**Supplementary Figure 2 | Loss of Rgy does not markedly alter Smc distribution at 85˚C**

**a** Genome-wide tracks of Smc ChIP-seq signal in KU216 (top panel) and Δ*rgy* cells (bottom panel). The cells were grown at 85˚C. Enrichment of immunoprecipitated versus input DNA (IP/input) is shown at 1-kb resolution.

**b, c** Smc ChIP-seq tracks around boundary 1 (**b**) and boundary 2 (**c**) were generated at 1-kb resolution. Solid blue lines: KU216 cells grown at 85˚C. Dashed orange lines: Δ*rgy* cells grown at 85˚C.

**d** Venn diagram showing the overlap of Smc ChIP-seq peaks between KU216 and Δ*rgy* cells grown at 85˚C.

**e** Smc IP/input values at the common Smc peaks between KU216 and Δ*rgy* cells grown at 85˚C.

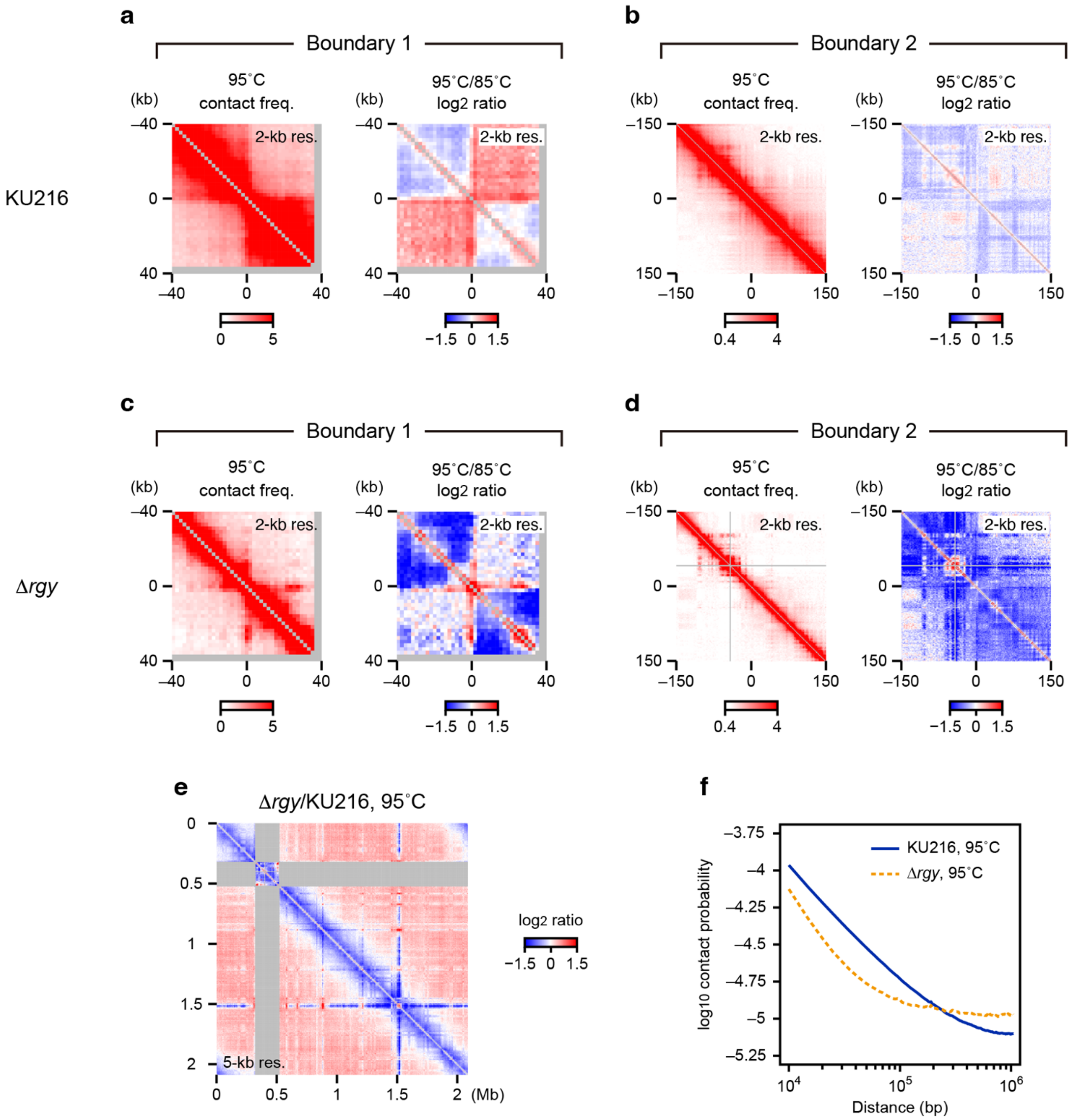


**Supplementary Figure 3 | Heat-induced 3D genome changes in Δ*rgy* cells**

**a, b** 3C-seq analysis at 2-kb resolution. KU216 cells were grown at 85˚C and treated at 95˚C for 1 h. Left panels: 3C-seq contact maps around boundary 1 (**a**) and boundary 2 (**b**). Right panels: differential contact maps showing $\log_2$ ratios of contact frequencies before and after the heat shock around boundary 1 (**a**) and boundary 2 (**b**).

**c, d** The same analyses as in **a** and **b** were performed on Δ*rgy* cells.

**e** Differential contact map visualizing $\log_2$ ratios of contact frequencies shown in Fig. 2a and b. The genomic contacts between the inverted region and the other loci (shaded in gray) were omitted from the analysis.

**f** Contact probability was plotted using 3C-seq data at 5-kb resolution. Solid blue line: KU216 cells treated at 95˚C for 1 h. Dashed orange line: Δ*rgy* cells treated at 95˚C for 1 h.

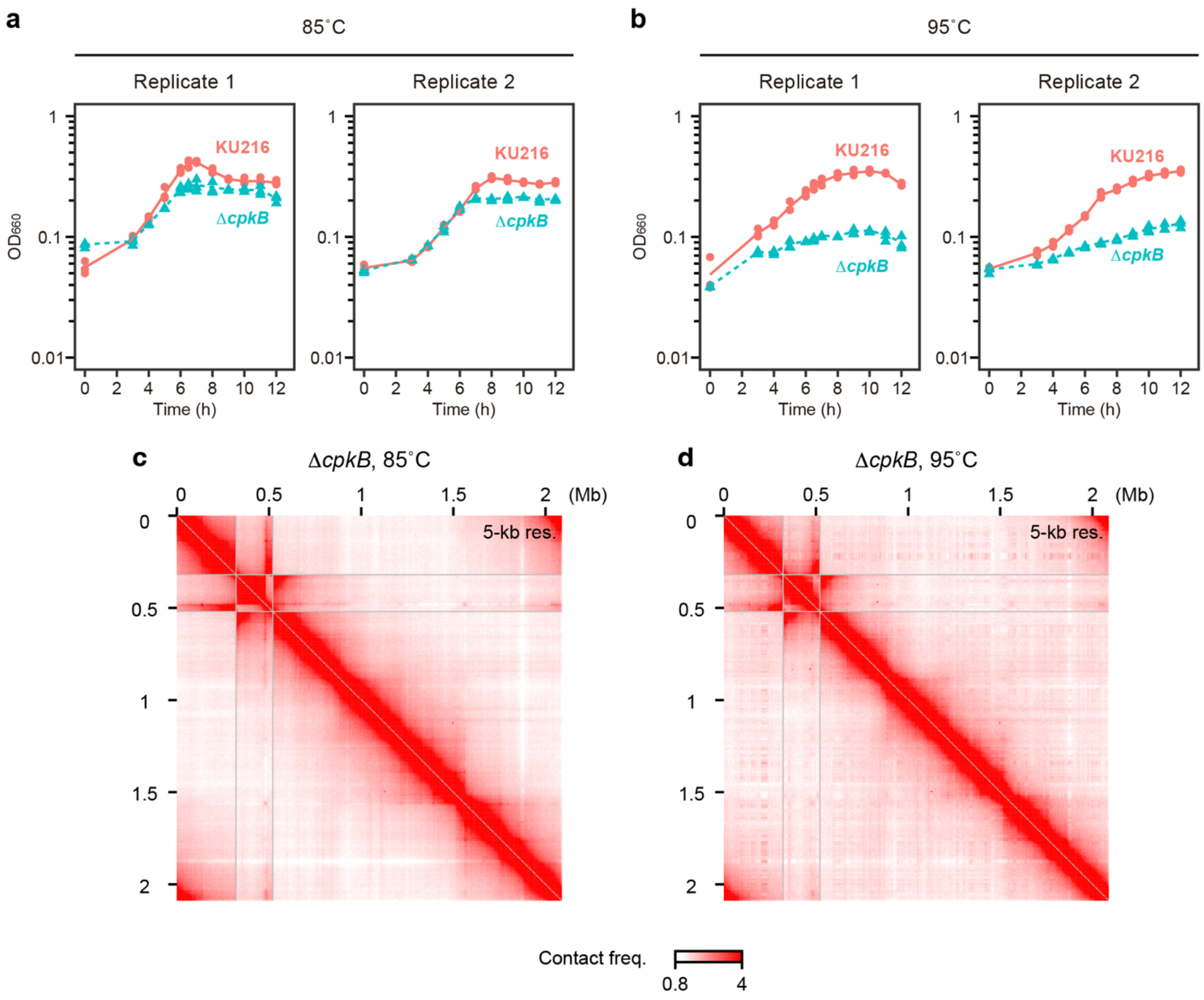


**Supplementary Figure 4 | Loss of the CpkB chaperonin impairs cell growth upon heat shock but does not result in heat-induced gene clustering**

**a, b** Growth curves of KU216 and a derivative Δ*cpkB* mutant were measured as in Fig. 4c. KU216: solid red circles and solid red lines. Δ*cpkB*: solid green triangles and dashed green lines.

**c, d** 3C-seq contact maps at 5-kb resolution. Δ*cpkB* cells were grown at 85°C (**c**) and exposed to 95°C for 1 h (**d**).

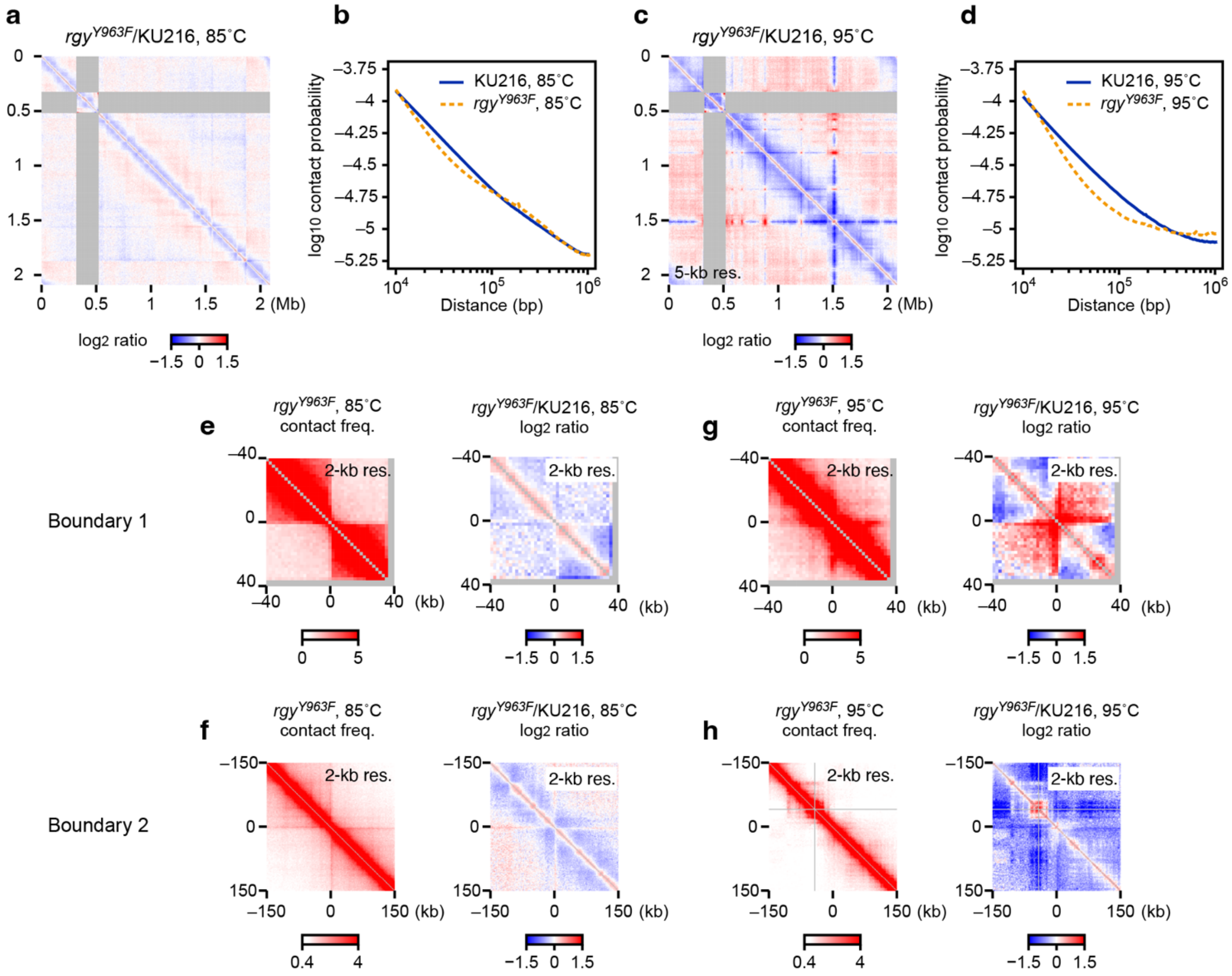


**Supplementary Figure 5 | *rgy*$^{Y963F}$ alters the genome architecture in a similar manner as Δ*rgy***

**a** Differential contact map visualizing $\log_2$ ratios of contact frequencies between KU216 and *rgy*$^{Y963F}$ cells grown at 85˚C. The genomic contacts between the inverted region and the other loci (shaded in gray) were omitted from the analysis.

**b** Contact probability was plotted using 3C-seq data at 5-kb resolution. Solid blue line: KU216 cells grown at 85˚C. Dashed orange line: *rgy*$^{Y963F}$ cells grown at 85˚C.

**c** Differential contact map as in **a** to visualize $\log_2$ ratios of contact frequencies between KU216 and *rgy*$^{Y963F}$ cells after the treatment with 95˚C for 1 h.

**d** Contact probability plot as in **b**. Solid blue line: KU216 cells treated at 95˚C for 1 h. Dashed orange line: *rgy*$^{Y963F}$ cells treated at 95˚C for 1 h.

**e, f** 3C-seq analysis at 2-kb resolution. KU216 and *rgy*$^{Y963F}$ cells were grown at 85˚C. Left panels: 3C-seq contact maps around boundary 1 (**e**) and boundary 2 (**f**). Right panels: differential contact maps showing $\log_2$ ratios of contact frequencies around boundary 1 (**e**) and boundary 2 (**f**).

**g, h** The same analyses as in **e** and **f** for KU216 and *rgy*$^{Y963F}$ cells treated at 95˚C for 1 h.

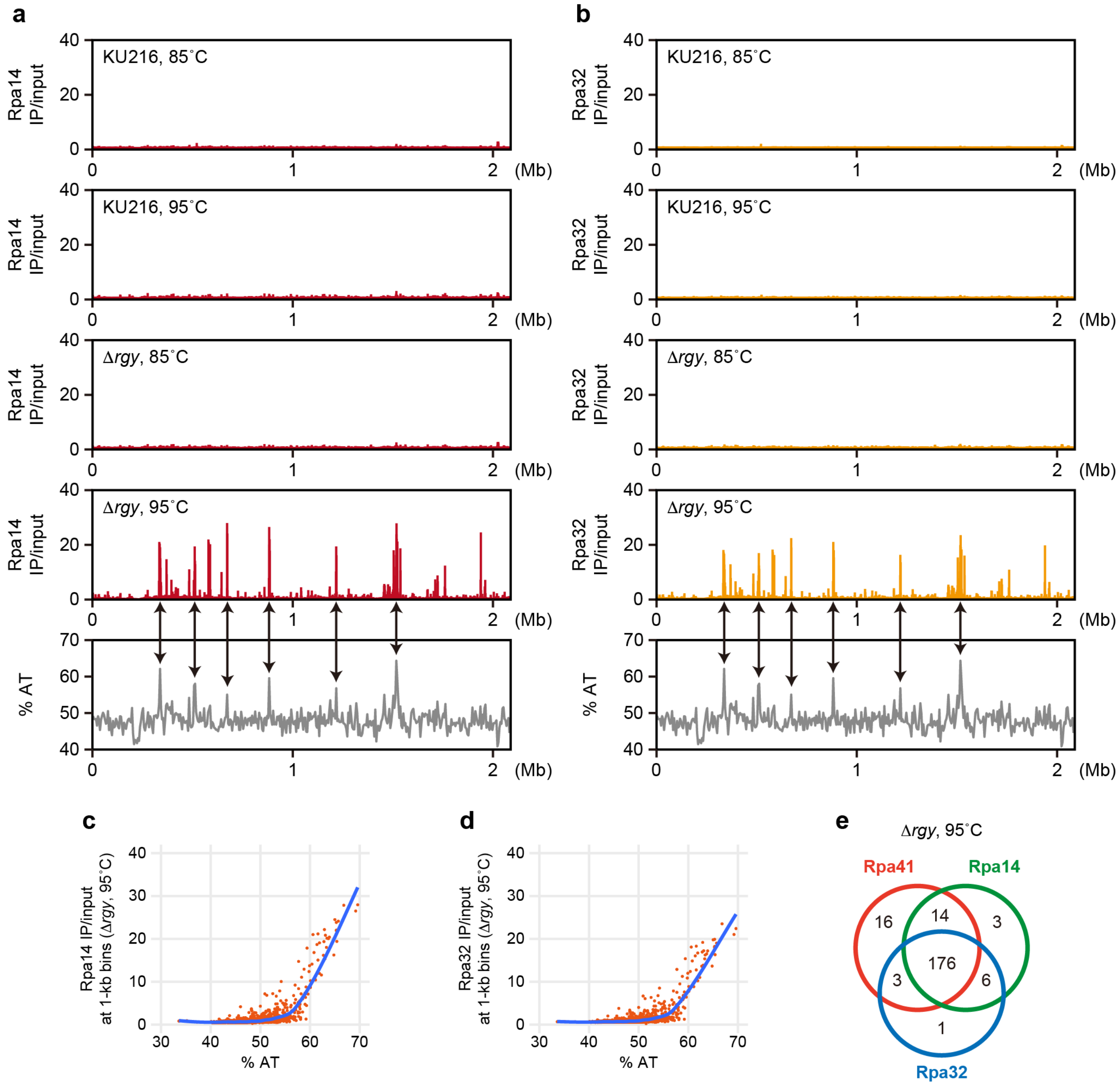


**Supplementary Figure 6 | Rpa14 and Rpa32 are enriched at AT-rich regions in heat-shocked Δ*rgy* cells**

**a** Top four panels: ChIP-seq tracks of Rpa14. KU216 and Δ*rgy* cells were grown at 85˚C (first and third from the top) and treated at 95˚C for 1 h (second and fourth from the top). The IP/input value is shown at 1-kb resolution. Bottom: AT-content track at 5-kb resolution. Overlaps between high Rpa14 peaks and AT-rich regions are indicated by arrows.

**b** Same analyses as in **a** for Rpa32.

**c** AT content and Rpa14 IP/input values were plotted for each 1-kb bin. Blue line: a LOESS regression curve generated with a span of 0.2.

**d** AT content and Rpa32 IP/input values were plotted as in **c**.

**e** Venn diagram showing the overlap of Rpa14, Rpa32, and Rpa41 ChIP-seq peaks in Δ*rgy* cells treated at 95˚C for 1 h.

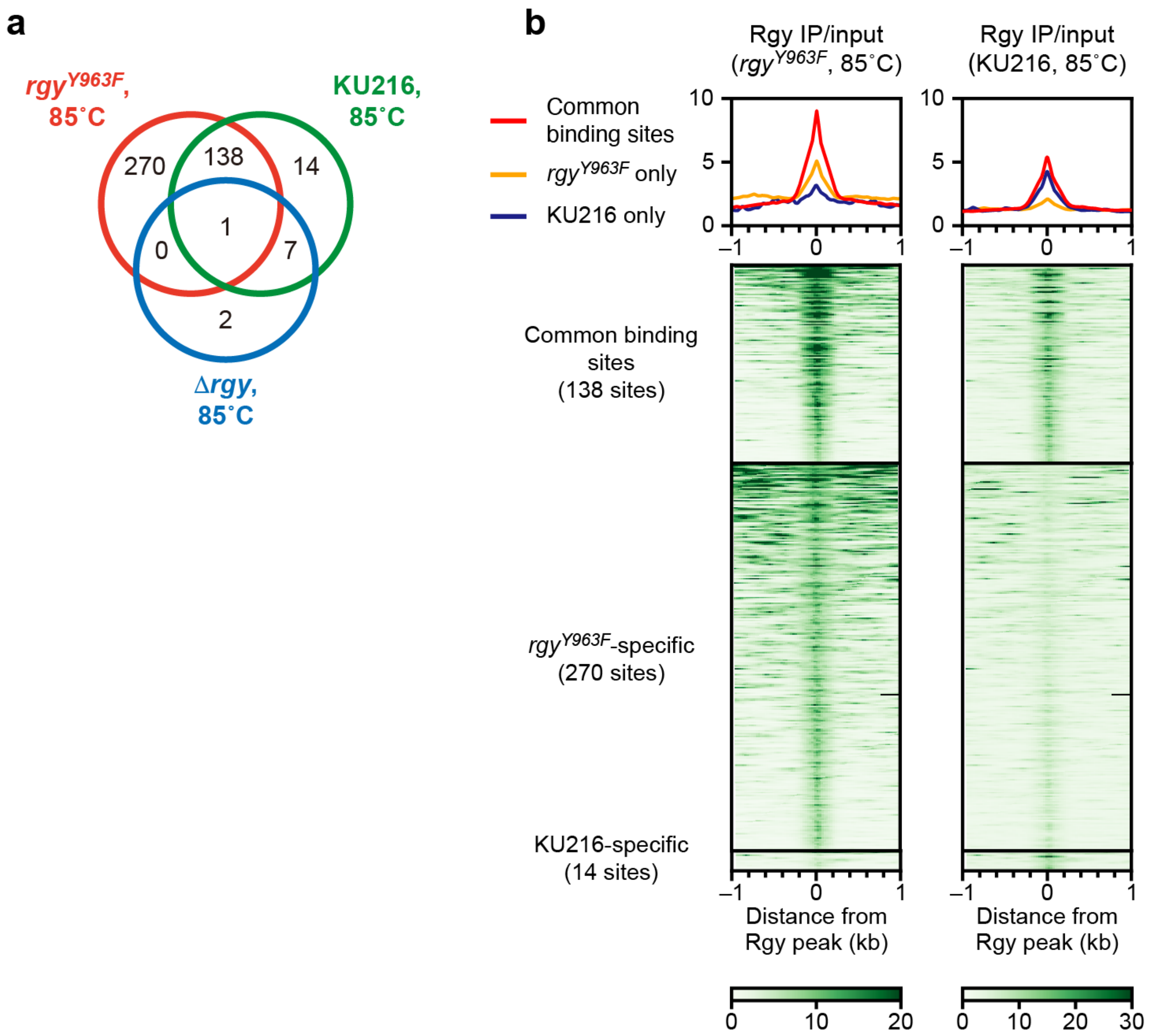


**Supplementary Figure 7 | *rgy*$^{Y963F}$ enhances Rgy binding to DNA**

**a** Venn diagram showing the overlap of Rgy ChIP-seq peaks identified in KU216, Δ*rgy*, and *rgy*$^{Y963F}$ cells grown at 85˚C.

**b** Average profiles (top) and heatmaps (bottom) of Rgy ChIP-seq signal were generated at 10-bp resolution. KU216 and *rgy*$^{Y963F}$ cells were grown at 85˚C. Rgy peaks found in the two groups were classified as common binding sites, *rgy*$^{Y963F}$-specific binding sites, and KU216-specific binding sites. Note that the Rgy peaks overlapping with those in Δ*rgy* cells were omitted as false positives.

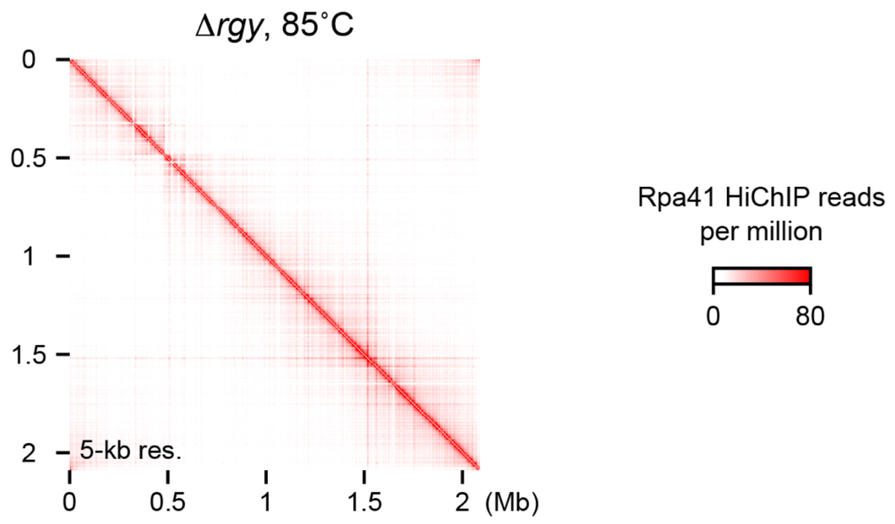


**Supplementary Figure 8 | RPA-bound loci do not form a spatial cluster in the absence of heat stress**

Rpa41 HiChIP contact map 5-kb resolution. Δ*rgy* cells were grown at 85˚C.

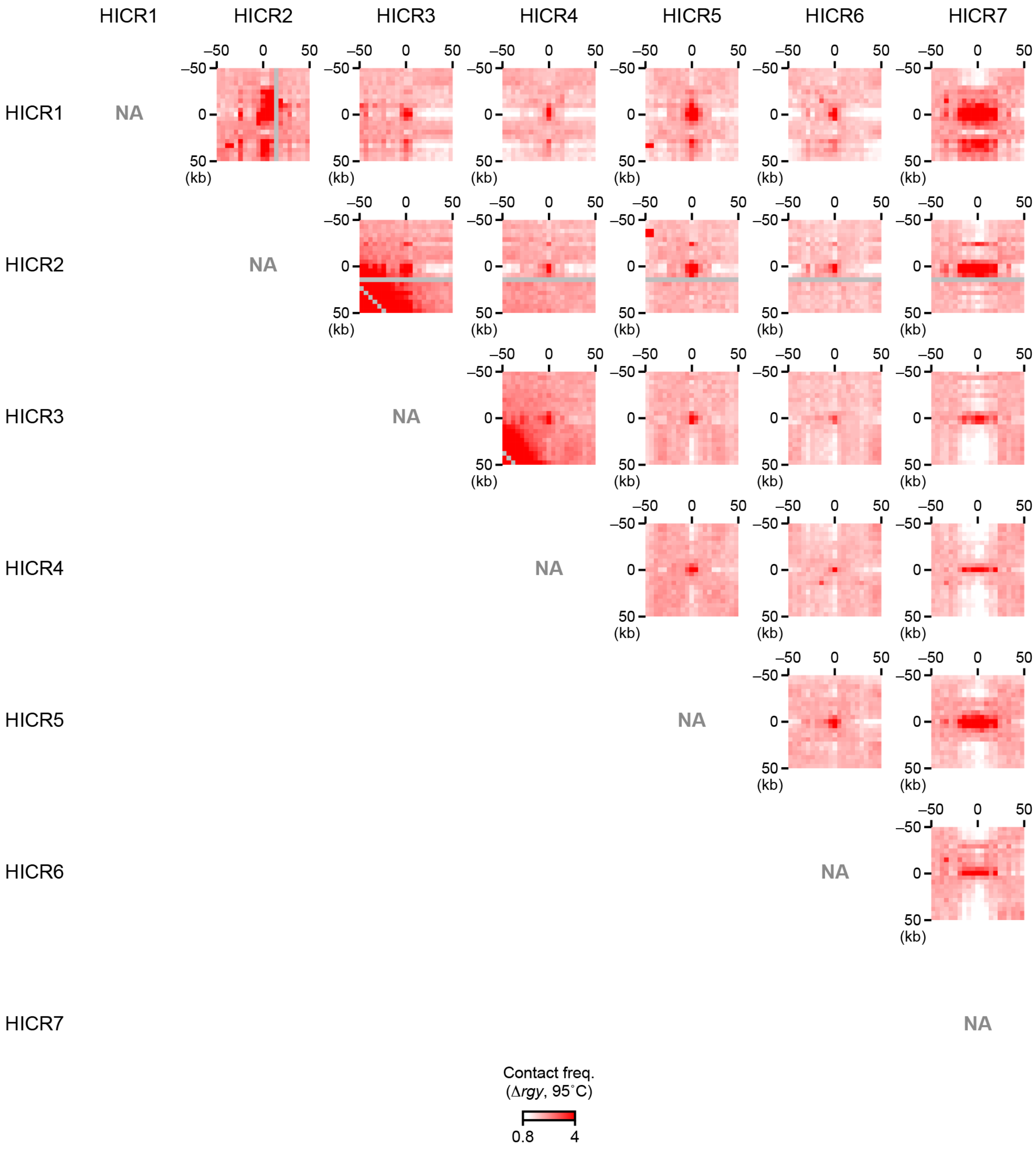


**Supplementary Figure 9 | The seven HICRs strongly interact with each other in heat-stressed Δ*rgy* cells**

Magnified views of the 3C-seq contact map in Fig. 2b were generated at 5-kb resolution, showing interactions among the ±50-kb regions centered on the HICRs.

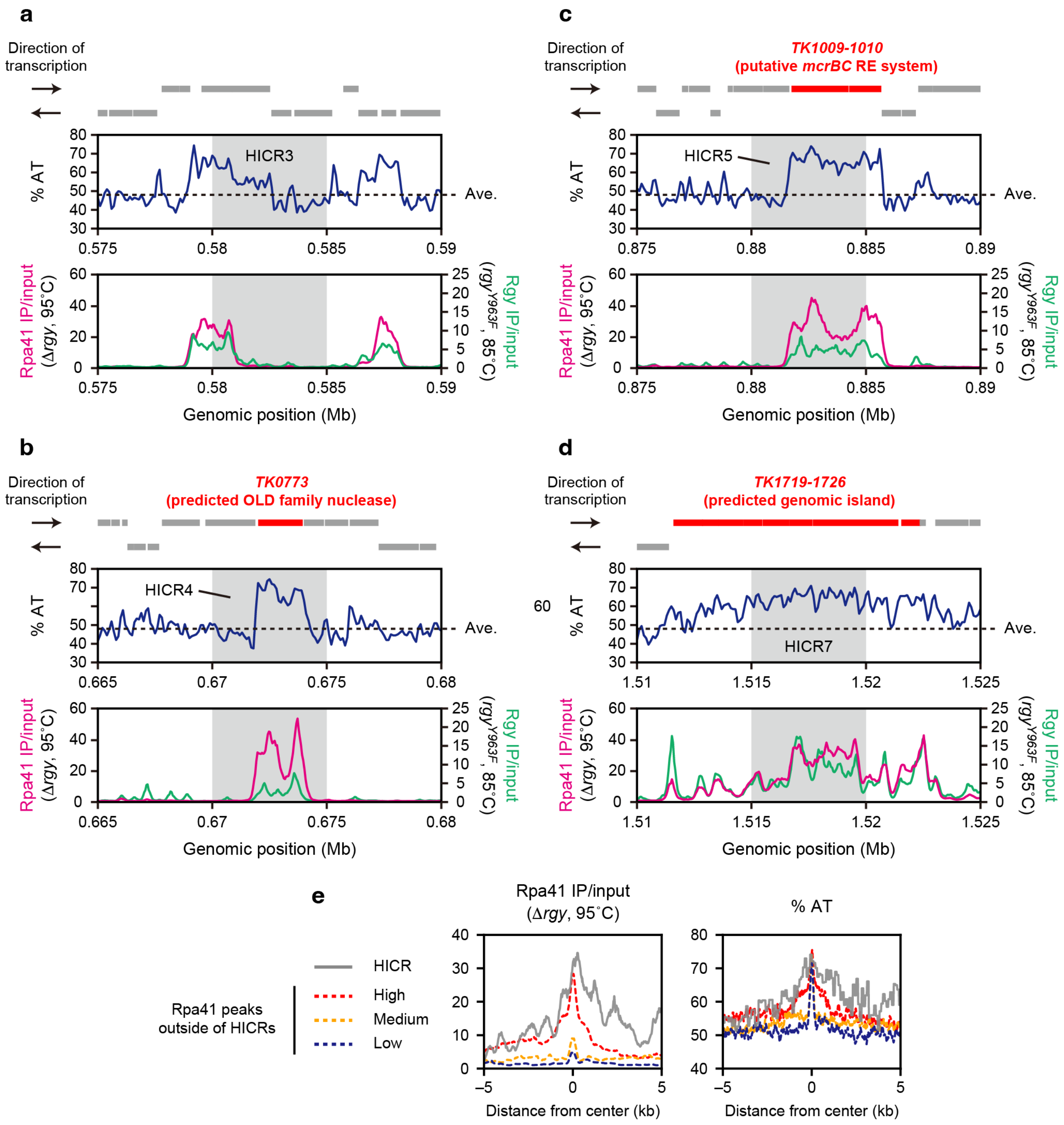


**Supplementary Figure 10 | Genomic characterization of HICRs**

**a-d** Genomic features of HICR3 (**a**), HICR4 (**b**), HICR5 (**c**), and HICR7 (**d**). Top panels: annotated genes (shown by gray lines) in the vicinity of the HICRs and their transcriptional directionality. Genes described in the main text are highlighted in red. Middle panels: AT-content tracks were generated with a sliding window size of 200 bp and a step size of 100 bp. The average AT content of the genome is shown by dashed horizontal lines. Bottom panels: an Rpa41 ChIP-seq track in Δ*rgy* cells treated at 95˚C for 1 h (magenta) and an Rgy ChIP-seq track in *rgy*$^{Y963F}$ cells grown at 85˚C (green) were generated at 50-bp resolution. In the middle and bottom panels, the locations of the HICRs are shaded in gray.

**e** Left panel: average profiles of Rgy ChIP-seq signal were generated at 10-bp resolution to compare Rgy enrichment at the HICRs (solid gray line) with those at the Rpa41 peaks outside of them. The latter loci were grouped based on the Rpa41 enrichment level. Right panel: average profiles of AT content were generated as in the left panel.

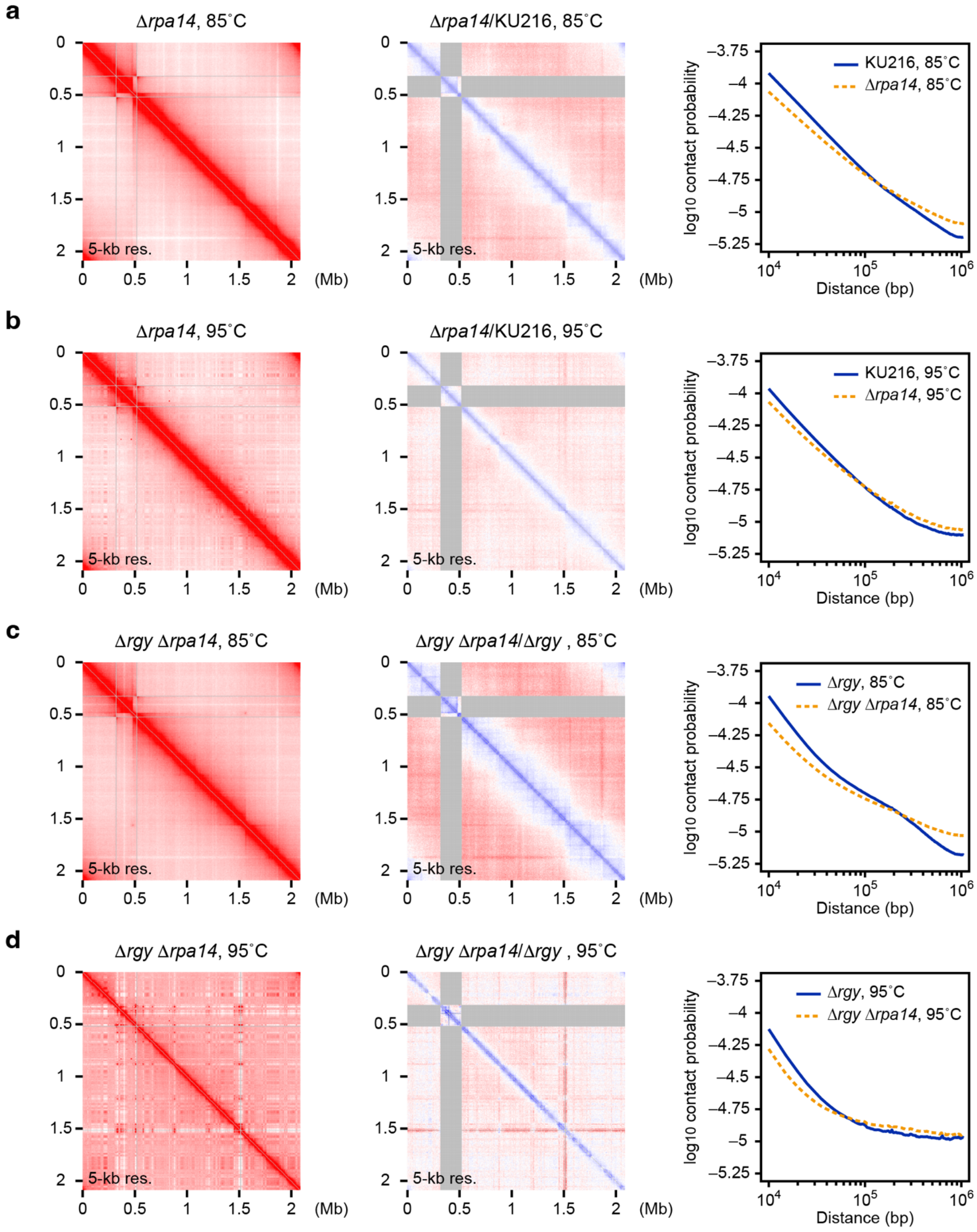


**Supplementary Figure 11 | Impacts of *rpa14* deletion on the 3D genome**

**a** Left panel: a 3C-seq contact map was generated at 5-kb resolution for Δ*rpa14* cells grown at 85˚C. Middle panel: differential contact map comparing contact frequencies in the left panel with those in KU216 cells grown at 85˚C (shown in Fig. 1a). The genomic contacts between the inverted region and the other loci (shaded in gray) were omitted from the analysis. Right panel: contact probability was plotted using 3C-seq data at 5-kb resolution. Solid blue line: KU216 cells grown at 85˚C. Dashed orange line: Δ*rpa14* cells grown at 85˚C.

**b** The same analyses as in **a** were performed to visualize the genome conformation in Δ*rpa14* cells treated at 95˚C for 1 h, and to compare it with that in KU216 cells treated at 95˚C for 1 h. In the left panel, the solid blue line and the dashed orange line represent the KU216 and Δ*rpa14* cells, respectively.

**c** The same analyses as in **a** were performed to visualize the genome conformation in Δ*rgy* Δ*rpa14* cells grown at 85˚C, and to compare it with that in Δ*rgy* cells grown at 85˚C. In the left panel, the solid blue line and the dashed orange line represent the Δ*rgy* and Δ*rgy* Δ*rpa14* cells, respectively.

**d** The same analyses as in **a** were performed to visualize the genome conformation in Δ*rgy* Δ*rpa14* cells treated at 95˚C for 1 h, and to compare it with that in Δ*rgy* cells treated at 95˚C for 1 h. In the left panel, the solid blue line and the dashed orange line represent the Δ*rgy* and Δ*rgy* Δ*rpa14* cells, respectively.
